# A scalable method to improve gray matter segmentation at ultra high field MRI

**DOI:** 10.1101/245738

**Authors:** Omer Faruk Gulban, Marian Schneider, Ingo Marquardt, Roy A.M. Haast, Federico De Martino

## Abstract

High-resolution (functional) magnetic resonance imaging (MRI) at ultra high magnetic fields (7 Tesla and above) enables researchers to study how anatomical and functional properties change within the cortical ribbon, along surfaces and across cortical depths. These studies require an accurate delineation of the gray matter ribbon, which often suffers from inclusion of blood vessels, dura mater and other non-brain tissue. Residual segmentation errors are commonly corrected by browsing the data slice-by-slice and manually changing labels. This task becomes increasingly laborious and prone to error at higher resolutions since both work and error scale with the number of voxels. Here we show that many mislabeled, non-brain voxels can be corrected more efficiently and semi-automatically by representing three-dimensional anatomical images using two-dimensional histograms. We propose both a uni-modal (based on first spatial derivative) and multi-modal (based on compositional data analysis) approach to this representation and quantify the benefits in 7 Tesla MRI data of nine volunteers. We present an openly accessible Python implementation of these approaches and demonstrate that editing cortical segmentations using two-dimensional histogram representations as an additional post-processing step aids existing algorithms and yields improved gray matter borders. By making our data and corresponding expert (ground truth) segmentations openly available, we facilitate future efforts to develop and test segmentation algorithms on this challenging type of data.

## Introduction

Magnetic resonance imaging (MRI) has become one of the most important tools to study human brain function and structure in vivo. Moving from high (3 Tesla [T]) to ultra high (7 and 9.4 T) magnetic fields (UHF), together with improvements in acquisition methods, leads to increases in signal and contrast to noise (SNR and CNR, respectively) [1–3]. The increase in SNR can be leveraged to increase the voxels’ resolution of both functional and structural images to sub-millimeter scales. Such sub-millimeter spatial resolutions allow for in vivo studies that probe cortical properties at the mesoscale [4–8]. These studies include (i) cortical-depth dependent analyses of function [9–14] and structure [5, 8, 15], (ii) the mapping of cortical columnar structures [16–21] as well as (iii) sub-millimeter cortical topography [18, 22–24].

Such studies crucially depend on accurate and precise delineations of the gray matter (GM) ribbon both at the inner (white matter; WM) and outer (cerebrospinal; CSF) border. Since the aim of these studies is to investigate how the (functional) MRI (f/MRI) signal varies as a function of small position changes in GM, systematic GM segmentation errors would invalidate the conclusions drawn from these studies. Consider, as an example, an fMRI study conducted with a voxel resolution of 0.8 mm isotropic. Assume an average thickness of human cortex of 2.4 mm and a true signal change at the upper cortical depth level. In this study, under optimal conditions the functional resolution would allow a straight piece of cortical ribbon to be divided in three relative cortical depths, each of them one voxel thick. Falsely labeling an additional fourth voxel as GM would make the difference between reporting an fMRI signal change at most superficial (false voxel excluded) or mid-superficial (false voxel included) cortical depth level. This example stresses the importance of accurate and precise GM segmentations.

Obtaining accurate and precise definitions of the GM ribbon, however, is currently a difficult and time-consuming task for sub-millimeter UHF data. The increases in SNR, CNR and resolution attainable in UHF anatomical data as well as analysis [25] and reconstruction [26] strategies specific to UHF reveal several structures outside of the brain that were barely visible on images obtained at conventional field strengths (1.5 and 3 T) and lower resolution (> 1 mm isotropic) [27]. Such structures include the dura mater [28], medium-sized blood vessels in the sulci [29] as well as draining sinuses and connective tissue adjacent to GM [30]. To date, many of the existing brain segmentation algorithms have been developed and benchmarked on images collected at 1 mm isotropic resolution or lower and at conventional field strengths [31] (but see [32]). If segmentation algorithms do not model these non-brain structures they might falsely label (part of) these structures as GM. Faced with such segmentation errors, researchers commonly correct the misclassified voxels manually. However, the increase in resolution leads to an exponential increase in the number of voxels, which renders manual correction a laborious task. Furthermore, manual correction is prone to error and may introduce an observer bias, thereby reducing the reproducibility of subsequent analyses [33]. This currently leaves researchers with the dilemma of accepting the likely erroneous outcome of automatic segmentation algorithms or performing a time-consuming and error-prone manual correction.

CBS tools [32] directly tackle many of the challenges of UHF high-resolution anatomical data by, for example, including pre-processing steps to estimate dura mater and CSF partial voluming. Consequently, these tools provide an improved initial GM segmentation compared to other solutions [32]. However, we show that in many cases the initial CBS segmentation can be further improved with the approaches proposed here. Furthermore, CBS tools have been optimized for whole-brain data obtained with the MP2RAGE sequence [26]. While the MP2RAGE sequence is commonly used at UHF as a basis for brain tissue class segmentations, we note that many high-resolution studies at UHF also use alternative sequences to define GM [12, 25, 34, 35], some of which offering partial coverage of the brain only [12, 35]. In such cases, alternative approaches that do not depend on particular templates, atlases or other forms of prior information are useful and required.

Here, we show that non-brain voxels misclassified as GM can largely be corrected using a multi-dimensional transfer function that is specified based on a two-dimensional (2D) histogram representation [36–39] of three-dimensional (3D) MRI brain data. We demonstrate that this transfer function offers an efficient way to single out non-brain tissue voxels. Removing these voxels from GM classifications found by automatic segmentation pipelines improves GM segmentations. This approach addresses the problems of an entirely manual correction, since it yields a meaningful summary representation of the data that allows to manipulate the data efficiently. As a consequence, it is both more time efficient than manual slice-by-slice correction and it reduces observer bias.

This paper is intended as a demonstration that 2D histogram based methods are useful for improving segmentation of MRI images. In particular, we aim to show that for images acquired at sub-millimetre resolution and at very high field strength (7 T and above) 2D histogram based methods offer an efficient way to obtain a more refined brain mask that excludes usually undesired structures like vessels and dura mater. Several alternatives to the methods presented here exist for data visualization, dimensionality reduction or data fusion, such as principal component analysis [40], multidimensional scaling [41] or the t-SNE algorithm [42]. However, a detailed quantification of the merits and disadvantages of these methods is beyond the scope of this manuscript which is intended to introduce a simple and fast solution. Likewise, there are alternative ways of defining clusters in an image to the normalized graph cut algorithm that we have used here [43–46]. While all these methods have their merit, we decided to use normalized-cut multilevel segmentation since it already has been shown to work successfully on the 2D histogram representations of volumetric data [39].

We structured the paper as follows. In Section 1, we introduce the technique of specifying transfer functions based on 2D histogram representations of voxel intensity and gradient magnitude. We offer theoretical considerations for why this technique is suited to remove vessels and dura mater voxels in high-resolution MRI data (< 1 mm isotropic voxel size). In Section 2, we extend the use of histogram-based transfer functions to multi-modal MRI data sets (e.g. T1 weighted [T1w], Proton Density weighted [PDw], T2* weighted [T2*w]) by considering MRI data in the compositional data analysis framework [47]. We show that this compositional framework yields an intuitive and useful summary representation of multi-modal MRI data which aids the creation of transfer functions. In Section 3 we outline required features of the input data and recommended data pre-processing steps. In Section 4 and 5 we validate the suggested methods by evaluating obtained GM segmentation results against expert GM segmentations obtained for nine subjects recorded at 7 T. We demonstrate considerable improvement in segmentation performance metrics for the two methods suggested here. We have implemented the methods described here in a free and open Python software package [48]. The package as well as validation data, corresponding expert segmentations [49], and processing scripts [50] used to validate the proposed methods are all openly available (see Table 1 for links).

**Table 1.** Availability of validation data and code. Validation data and scripts as well as segmentation software are all openly accessible by following the corresponding links for their repositories.

| What? | Where? |
| --- | --- |
| data | https://zenodo.org/record/1206163 |
| scripts | https://zenodo.org/record/1219231 |
| software | https://zenodo.org/record/999487 |

## 1 Theory I: Transfer functions and 2D histograms

### 1.1 Multi-dimensional transfer functions

In the context of MRI data visualization, a transfer function can be understood as a mapping of voxel data to optical properties such as color and opacity. Effective transfer functions make structures of interest in the data visible. This can, for example, be achieved by assigning low opacity values to voxels that make up irrelevant structures and by highlighting desired structures with high opacity and salient color values. Multi-dimensional transfer functions assign renderable properties based on a combination of values [36–38, 51]. This is important in the context of MRI data where features of interest are often difficult to extract based on a single value alone. Considering multiple values, such as the intensity in images acquired using different contrast weighting (e.g. T1w, PDw and T2*w), increases the chances of uniquely isolating a feature and making it visible [51].

In theory, multi-dimensional transfer functions could be used to perform exhaustive tissue-type segmentation of human brain MRI data. In this process, each voxel would be classified as either WM, GM, or CSF by specifying appropriate transfer functions. It has been shown, however, that this approach is less successful than other, bespoke brain segmentation algorithms [52]. Here, we propose that transfer function-based methods still have a role to play in UHF MRI brain segmentation pipelines because they are well-suited for efficient removal of mislabeled non-brain tissue. We motivate this proposition by considering that brain and non-brain voxels become separable in 2D histogram representations.

### 1.2 2D histogram representation

2D histogram representations have been shown to greatly facilitate the process of specifying effective, 2D transfer functions [36, 38]. 2D histograms are obtained by taking an n-dimensional data set and binning its data points along two dimensions. In principle, a 2D histogram can be obtained from any two sets of values. A 2D histogram, plotting gradient magnitude against image intensity, however, has been shown to be particularly useful to identify tissue boundaries [36, 38]. The term gradient magnitude here refers to the magnitude of the vector that represents the spatial intensity gradient at every MRI voxel, where the spatial intensity gradient is equal to the first spatial derivative of the image intensity values.

Fig 1 shows how 3D MRI data of a human brain are represented in a 2D histogram (a T1w image was divided by a PDw image [25] and brain extracted; images were acquired with 0.7 mm isotropic resolution; for more details see Section 4). The histogram is obtained by plotting gradient magnitude against image intensity. In this representation, different tissue types occupy different regions. CSF voxels are characterized by very low intensity and low gradient magnitude values and therefore occupy the lower-left space of the histogram. GM and WM voxels have medium to high intensities and very low gradient magnitudes. Therefore, these tissue classes form circular regions at the bottom-center of the histogram. Voxels at the GM-WM interface fall within an arc reaching from medium to high intensities, following a low to medium to low gradient magnitude trajectory. Similarly, voxels at the CSF-GM interface span an arc from low to medium intensities. Finally, blood vessels and dura mater are thin structures characterized by very high gradient magnitudes and medium to high intensities. Therefore, these structures occupy up-center and the up-right parts of the 2D histogram.

**Fig 1.**
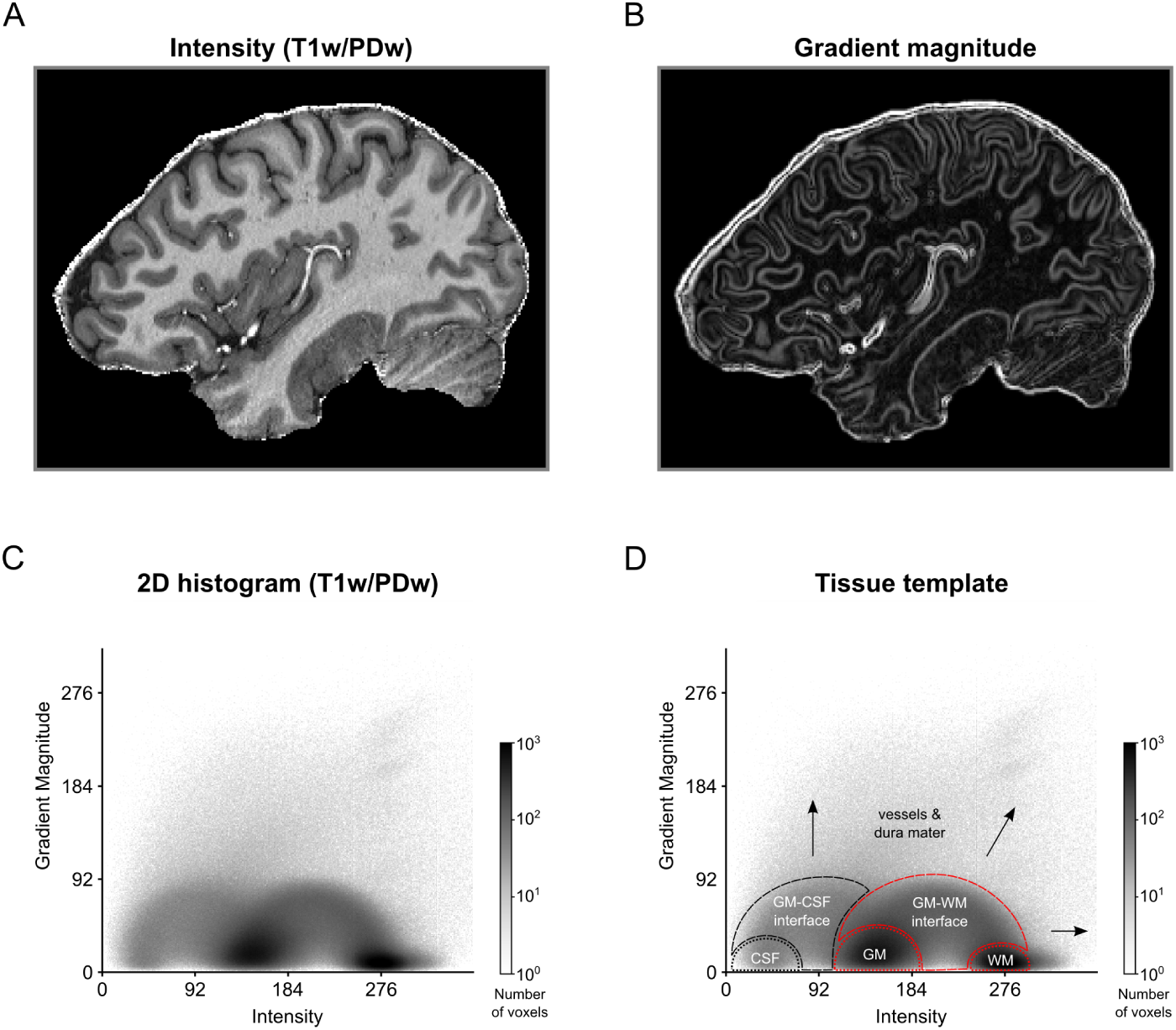
2D histogram representation for MRI image of a human brain. (A) Intensity and (B) gradient magnitude values of a brain extracted T1w-divided-by-PDw MRI image are represented in a (C) 2D histogram. Darker regions in the histogram indicate that many voxels in the MRI image are characterized by this particular combination of image intensity and gradient magnitude. (D) The 2D histogram displays a characteristic pattern with tissue types occupying particular areas of the histogram. Voxels containing CSF, dura mater or blood vessels (black dashed lines and arrows) cover different regions of the histogram than voxels containing WM and GM (red dashed lines). As a result, brain tissue becomes separable from non-brain tissue.

Since different tissue types occupy different regions in the 2D histogram, each tissue type and boundary can, in principle, be isolated using a 2D transfer function based on image intensity and gradient magnitude. For the purposes of this paper, we focus on the distinction between brain (WM, GM, GM-WM interface) and non-brain (CSF, CSF-GM interface, blood vessels, dura mater) voxels. The intensity-gradient magnitude histogram is particularly suited to distinguish non-brain tissue because voxels containing dura mater and vessels are characterized by high gradient magnitude values. Given the typical voxel sizes of current high resolution studies, gradient magnitude will be high in the entirety of these structures (see Fig 1B for an example) and the combination of high intensity and high gradient magnitude values renders these structures separable from WM and GM voxels.

### 1.3 Creating transfer functions

The simplest way to create a transfer function is to explore the data by moving widgets with a specified shape over the 2D histogram representation [38]. For example, Fig 2 shows how a circular sector could be moved on top of the 2D histogram to highlight particular regions. In this case, only MRI voxels whose intensity-gradient magnitude combination falls within the highlighted region of the 2D histogram would be selected. Position and size of the circular sector can then be refined until the desired data have been isolated.

**Fig 2.**
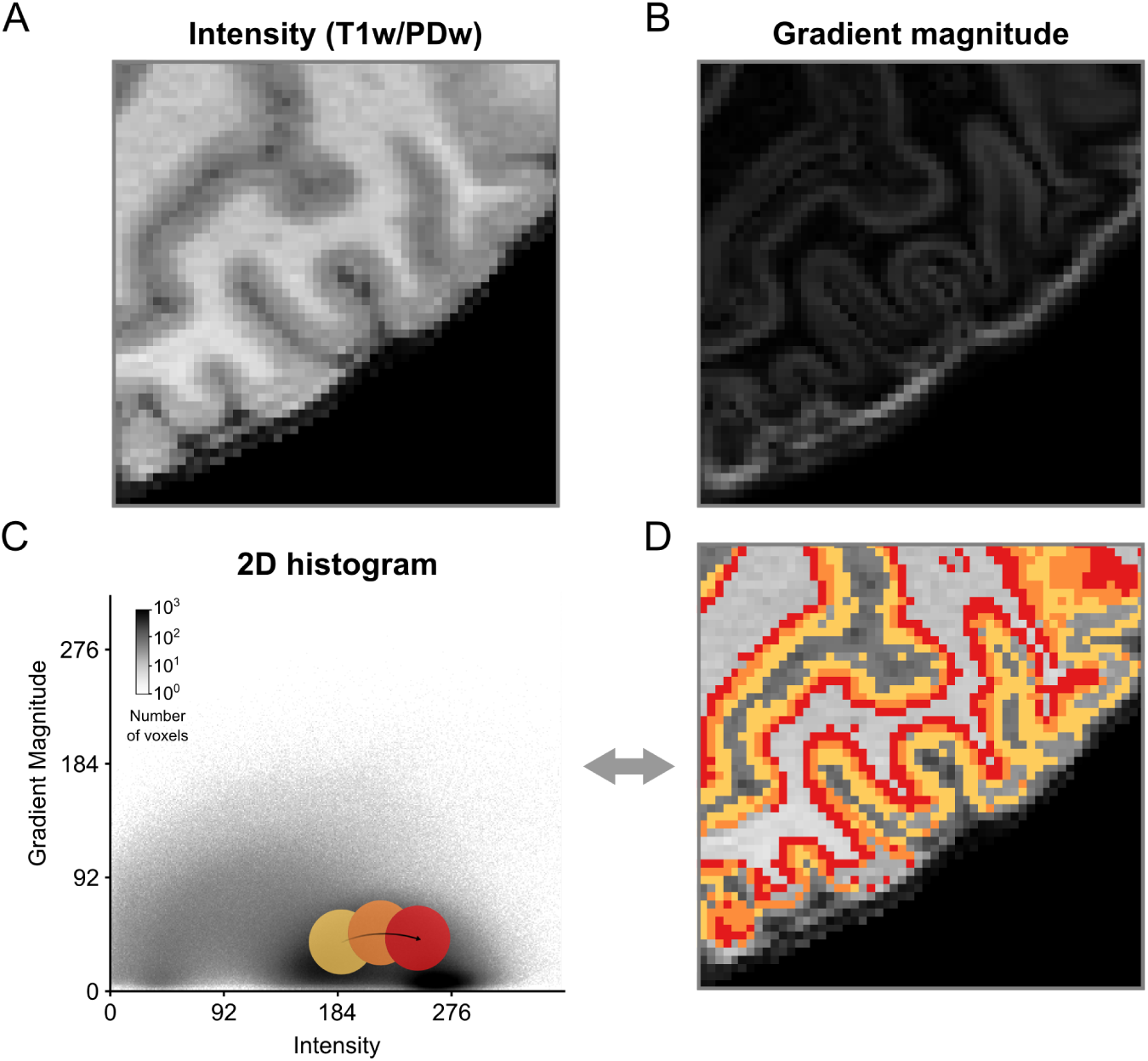
Creation of 2D transfer functions with pre-defined shapes. (A) Intensity and (B) gradient magnitude values of of a brain extracted T1w-divided-by-PDw MRI image are represented in a 2D histogram. By moving widgets of pre-defined shape, e.g. a circle, over the (C) 2D histogram and (D) concurrent visualization of selected voxels on a 2D slice of brain, positions of different tissue types in the 2D histogram can be probed and transfer functions can be created. In this example, the different probe positions (yellow, orange and red circles) appear to contain different aspects of GM.

Using such a straightforward process of exploration and refinement [51], however, might yield slightly sub-optimal results. The shape of the widget might not capture the ideal shape given the data or the user might lack the prior knowledge that is required for this task. Alternatively, hierarchical exploration of normalized graph cut decision trees [39] can be used. This graph cut method results in a set of components (i.e. clusters) of the histogram that are mutually exclusive and collectively exhaustive. This allows the user to split and merge clusters in a data-driven and intuitive way that can be aided by the immediate visualization of the resulting segmentation (Fig 3, S1 Video). The method allows for semi-automatic tissue selection, i.e. the shape of the clusters is data-driven but the decision which clusters to join and which to divide is made by the user.

**Fig 3.**
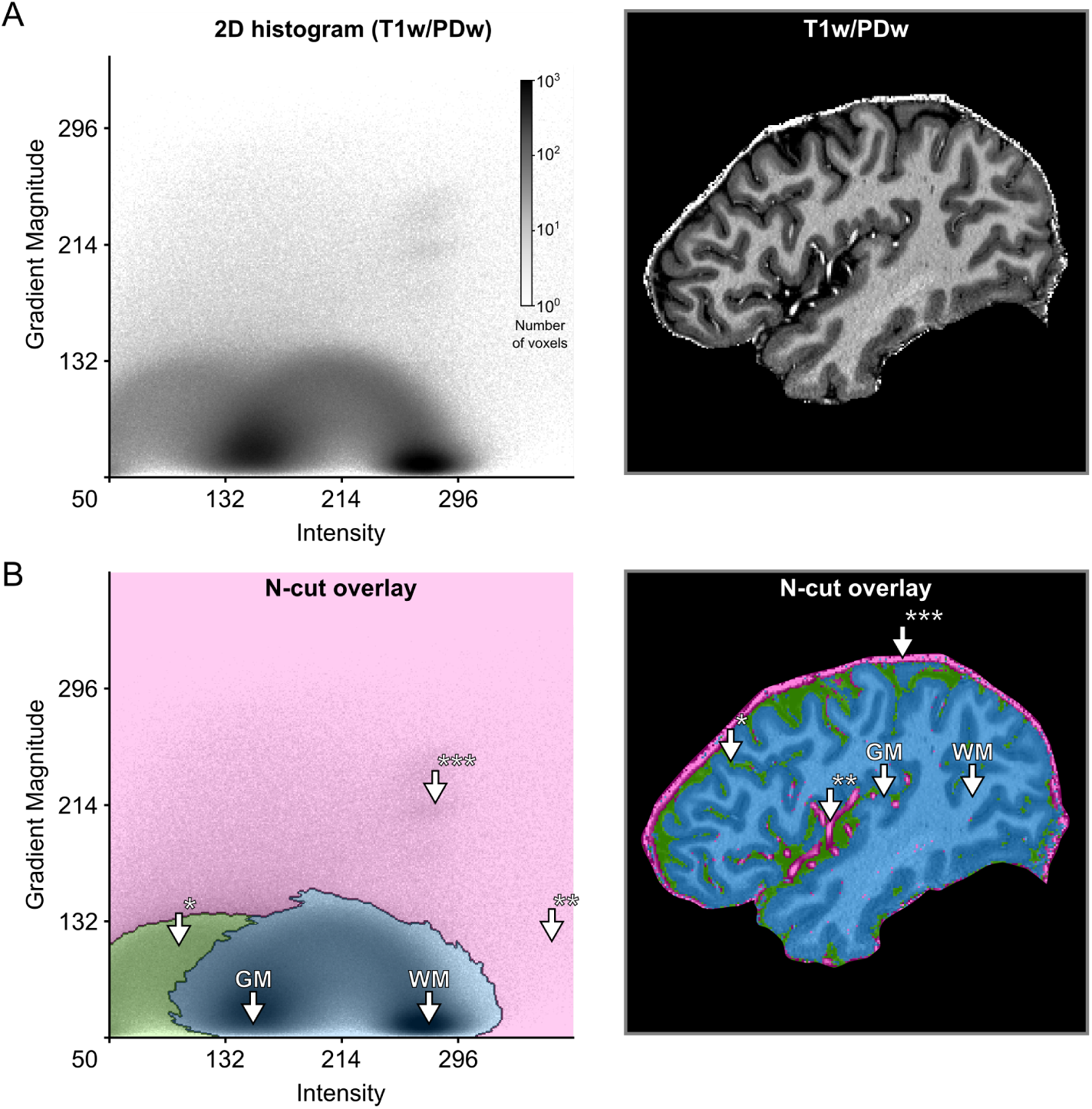
Creation of 2D transfer functions with data-driven shapes. (A) The user starts with the 2D histogram representation of image intensity and gradient magnitude (left side) and concurrent visualization of the original brain data (right side). The user can then interact with and select data in the 2D histogram to specify transfer functions. In this example, this was done with the help of a normalized graph cut decision tree. (B) The interaction with the 2D histogram results in data-driven shapes of selected areas, here shaded in pink, green and blue (left side). Voxels selected by those areas are highlighted in corresponding colors against the backdrop of the original brain data (right side). The visualization reveals that the area of the 2D histogram shaded in blue selects brain voxels, while the areas shaded in green and pink select CSF* and blood vessel voxels**/dura mater***, respectively.

## 2 Theory II: Multi-modal MRI data analysis

### 2.1 Compositional analysis for MRI data

More than one MRI contrast is often available and a combination of different contrasts can be useful in distinguishing different tissue types by differentially highlighting unique intrinsic properties. Two images with different contrast weighting can be combined using, for example, a ratio image [25, 53, 54]. This approach is beneficial for two reasons:1) it reduces image biases as all acquisitions are affected by the same sensitivity profile of the receive elements in the radio frequency (RF) coil, and 2) if the images carry opposing contrast for the tissues of interest, the ratio increases contrast and benefits the delineation of the structures (tissues) of interest.

The ratio image approach, however, is limited to pairs of images. To operate on the relative information of more than two images, we propose to use the barycentric coordinate system which was discovered by August Ferdinand Möbius in 1827 [55]. In the barycentric coordinate system, coordinates of a point represent a simplex whose center of mass is determined by the weights at its vertices (the term *n-simplex* in geometry is the generalized form of the triangle [56]; for example the 0-simplex is a point, the 1-simplex is a line segment, the 2-simplex is a triangle, the 3-simplex is a tetrahedron and so on). In other words, points in the barycentric coordinate system represent compositions of non-negative fractions whose sum of components gives a constant value. The barycentric coordinates of multiple measurements acquired in each voxel can be extracted through the following vector decomposition:

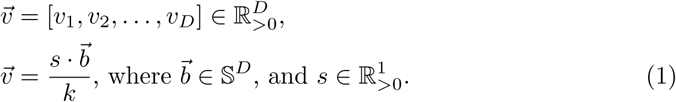

The vector 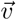 stands for a voxel with *D* number of measurements, 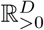 indicates positive real numbers, *k* is an arbitrary scalar, *s* is a scalar representing sum of the vector components and the vector 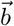 stands for the barycentric coordinates which belong to simplex sample space 𝕊^*D*^. The barycentric coordinates are acquired by applying closure operation (*C*) used in compositional data analysis [47] to 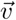:

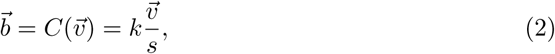

This decomposition (Eq. 1) might seem trivial, however the statement highlights the sampling space of the component 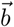 which is *D* dimensional simplex 𝕊^*D*^. When a set of measurements are represented as vectors with positive components summing up to a constant (e.g. percentages), compositional data (CoDa) analysis methods [47, 57, 58] becomes relevant. The compositional data analysis offers a set of principled operations taking the geometry of the simplex space into account. The general framework for CoDa analysis and its fundamental operations have already been rigorously documented in [47], however, for completeness we provide a step-by-step illustration of how multi-modal MRI data with three image contrasts (here T1w, PDw and T2*w magnitude images) can be processed under the compositional data analysis framework to acquire a useful representation of different tissue types. By only analyzing the barycentric components, the data are being compressed resulting in some information loss. However, this compression is done with the aim of revealing more useful information through the remaining components.

Let multi-modal MRI data consisting of T1w, PDw, T2*w measurements be defined as a matrix **M** with *n* rows and 3 columns (in relation to Eq. 1 *D* = 3):

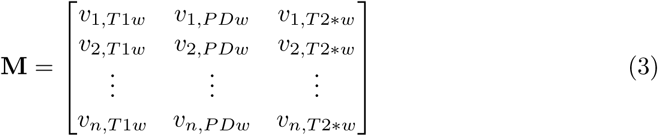

where *n* stands for the total number of voxels and each row *v_i_* is the vector of measurements for a specific voxel *i*. Each column represents an image.

The first step in compositional MRI data analysis is to convert the data components from Cartesian coordinates in real space (ℝ^3^) to barycentric coordinates in simplex space (𝕊^3^), applying the closure operation (Eq. 2) to every voxel (i.e. to each row of **M**) for obtaining a new matrix **B** indicating the barycentric coordinates of every voxel:

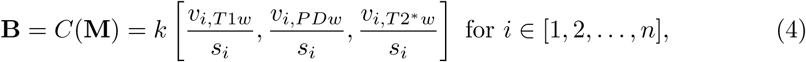

*k* can be ignored after selecting it as 1.

It is important to note that in the case of MRI images the scalar component *s* by itself does carry information; however, this information relates to the bias field in cases where the bias field is approximately equal across measurement types. Since we are not interested in bias field information, we do no longer use this component.

As the next step, the barycentric coordinates of compositions (**B**) are centered (i.e. normalized) by finding the *sample center* and *perturbing* each composition with the inverse of the sample center:

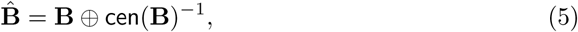

where the symbol ⊕ denotes the perturbation operation defined in multi-dimensional simplex space (*S^D^*), which can be considered as an analogue of addition in real space:

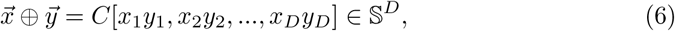

where 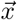 and 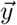 indicates two different compositions consisting of *D* components and `cen`(*B*) stands for:

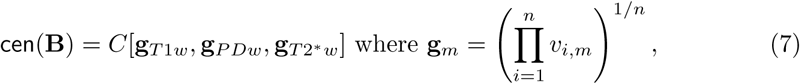

where *n* is the number of voxels, *C* is the closure operator (Eq. 2) and g*_m_* is the geometric mean of component *m* (i.e. T1w, PDw, T2*w).

After centering, the data are standardized:

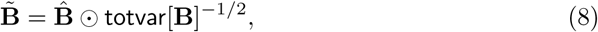

the symbol ☉ stands for the power operation defined in simplex space, which can be considered as an analogue of multiplication in real space:

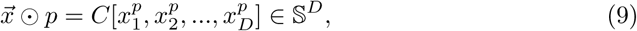

where 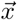 is the barycentric coordinates of a composition with *D* components and *p* is a scalar. The total variance in Eq. 8 is computed by:

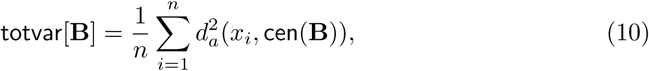

where 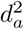 indicates squared Aitchison distance. This is a metric defined in simplex space that is analogous to Euclidean distance in real space:

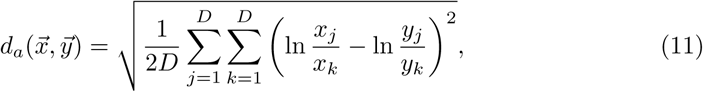

the barycentric coordinates 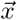 and 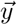 indicate two different compositions consisting of *D* components. For example in the case of compositions consisting of T1w, PDw and T2*w measurements *D* = 3.

After standardization, the barycentric coordinates are transformed from the three dimensional simplex space 𝕊^3^ to two dimensional real space (ℝ^2^) with the purpose of conveniently visualizing the compositional distribution in a 2D histogram by using the isometric logratio (`ilr`) transformation [59]:

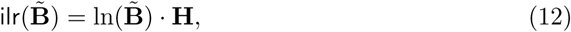

where `ilr` transformation is applied to every voxel and **H** indicates a Helmert sub-matrix [60] of 3 rows and 2 columns:

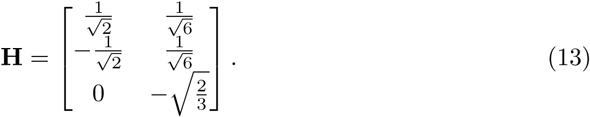

We have selected the matrix **H** because it is the suggested standard choice [61].

Note that the closure operation described in Eq. 2 implies scale invariance. If the receive (and in some cases transmit) field (B1) inhomogeneities for MRI data are similar across modalities and assumed to be having a multiplicative effect on the measured signal, applying closure will mitigate inhomogeneities by canceling out the common multiplicative term (ie. bias field) in each image modality. For instance, assume two voxels contain the same tissue type but have dissimilar intensities due to a multiplicative effect. If before the closure operation voxel 1 has an intensity of 100 in all recorded modalities and voxel 2 has an intensity of 500 in all modalities, then after the closure operation both voxels will have the same compositional description, which would be desired. It should be noted that if B1 inhomogeneities differ significantly across modalities, the closure operation will yield inaccurate compositional descriptions. In this case, we recommend to use bias field correction algorithms before using the compositional data analysis framework. A practical example for this case is that for magnetization-prepared rapid acquisition gradient-echo (MPRAGE) sequences, the transmit field in T1w image is effected by an inversion pulse which is not present in PDw and T2*w images. In such cases, individual image bias field correction is recommended.

### 2.2 2D histogram representation and creation of transfer functions

Fig 4 shows how three different 3D MRI contrast images of a human brain (T1w, PDw and T2*w brain extracted images; 0.7 mm isotropic resolution; for more details see Section 4) can be represented in a 2D histogram. The 2D histogram is obtained by taking the three MRI contrast images as an input and performing the operations of the CoDa analysis framework described above. In particular, applying the *ilr* transformation to the barycentric coordinates allows the three images to be represented along two dimensions. Different tissue types have different compositional characteristics and therefore occupy different regions in the resulting 2D histogram. WM and GM voxels are separated in two distinct clusters which mainly differ along the T1w axis. CSF voxels occupy the lower left corner of the histogram, which represents a combination of low T1w with high PDw and T2*w values. CSF voxels still differ from WM and GM voxels mainly along the T1w axis. In contrast, vessel and dura mater voxels differ from WM, GM and CSF voxels also along the PDw and T2*w axes, which makes these voxels to be spread out in the direction orthogonal to the T1w axis. To see how a combination of two MP2RAGE images (UNI, INV2) and one T2* image estimated from a multi-echo 3D gradient recalled echo (GRE) sequence are represented in a 2D histogram, please see S7 Fig.

**Fig 4.**
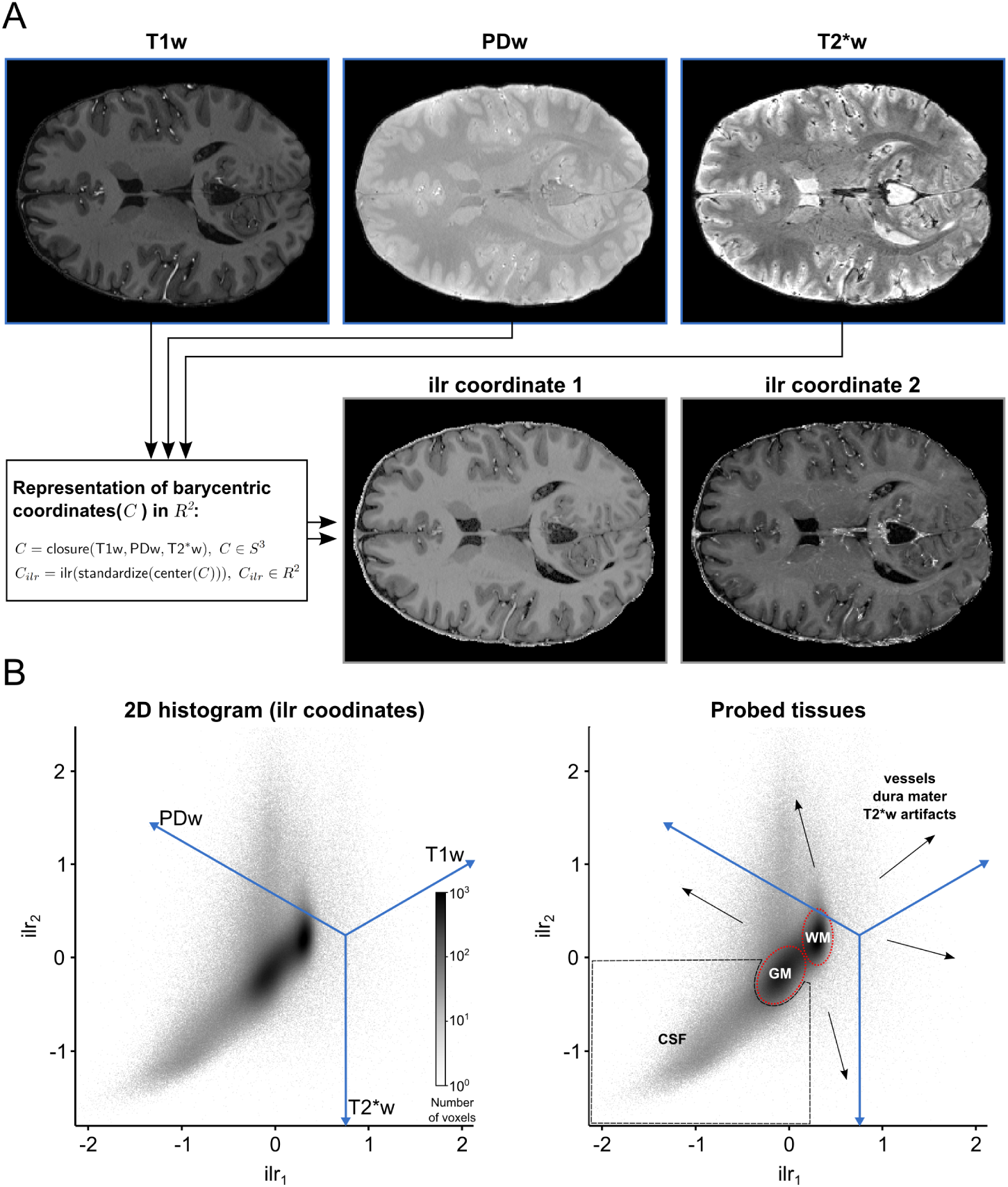
2D histogram representation of three 3D MRI contrast images. (A) Each voxel is considered as a 3 part composition in 3D real space. The barycentric coordinates of each composition which reside in 3D simplex space are represented in 2D real space after using a isometric log-ratio (ilr) transformation. (B) The ilr coordinates are used to create 2D histograms representing all voxels in the images. The blue lines are the embedded 3D real space primary axes. It should be noted that in this case the ilr coordinates are not easily interpretable by themselves but they are useful to visualize the barycentric coordinates which are interpretable via the embedded real space primary axes. Darker regions in the histogram indicate that many voxels are characterized by this particular scale invariant combination of the image contrasts. In this representation, brain tissue (WM and GM, red dashed lines) becomes separable from non-brain tissue (black dashed lines and arrows).

The dimensionality reduction accomplished by the *ilr* transformation allows to specify 2D transfer functions even though the input consists of three channels. Fig 5 shows how normalized graph cuts can be used on 2D histogram representation of *ilr* coordinates to create transfer functions. The resulting transfer functions highlight specific clusters that readily separate brain tissue from non-brain tissue.

**Fig 5.**
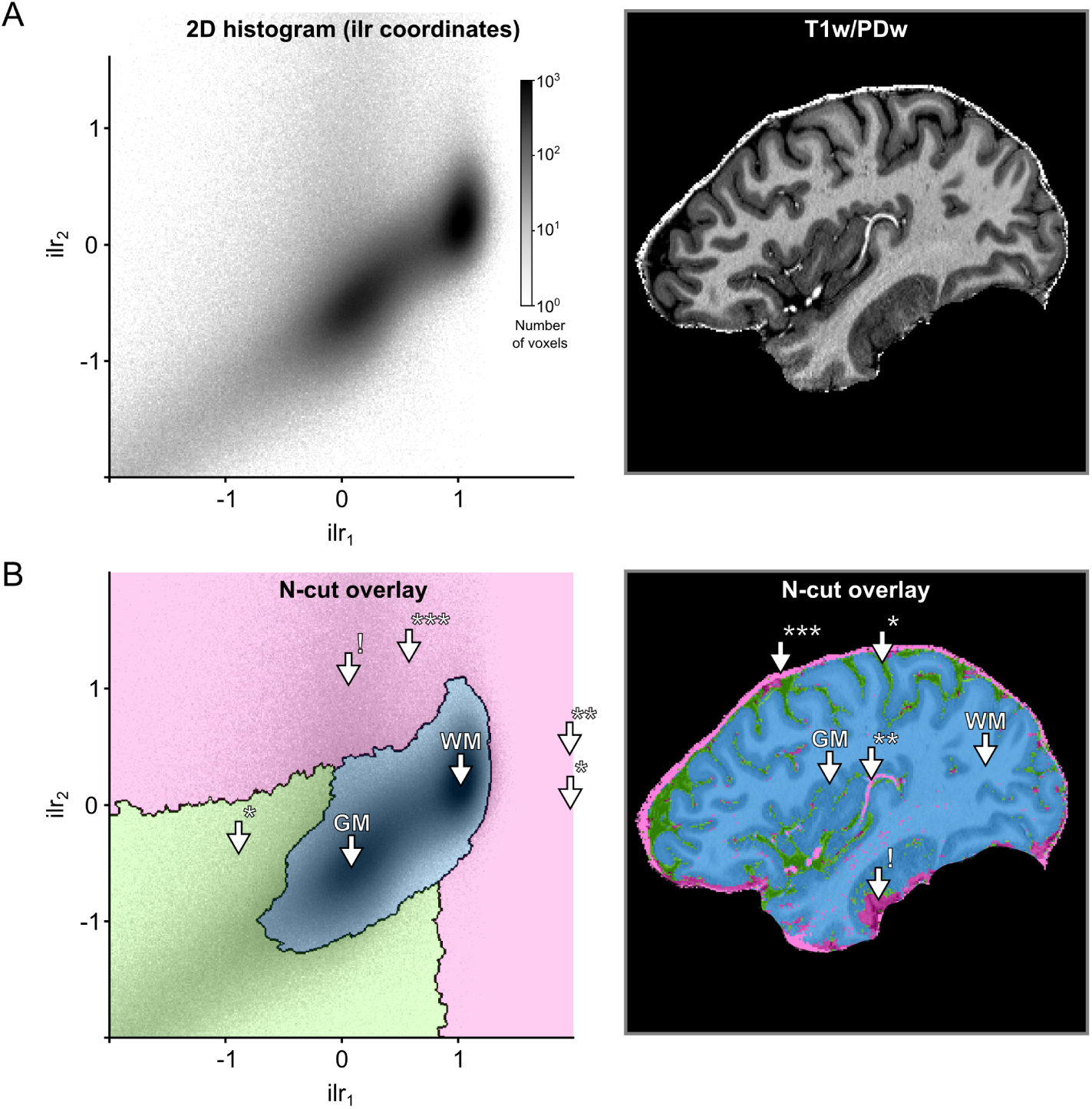
Creation of transfer functions using ilr coordinates. (A) The user starts with the 2D histogram representation of ilr coordinates 1 and 2 (left side) and concurrent visualization of the original brain data (right side). The user can then interact with and select data as described in Fig. 3. (B) The interaction with the 2D histogram results in data-driven shapes, here shaded in pink, green and blue (left side). Voxels selected by those areas are highlighted in corresponding colors against the backdrop of the original brain data (right side). The visualization reveals that the area of the 2D histogram shaded in blue selects brain voxels, while the areas shaded in green and pink select CSF* and blood vessel voxels**/dura mater***, respectively. The arrow with exclamation mark (!) indicates an area affected by T2*w image artifacts.

## 3 Input data requirements and preparation

### 3.1 Data preparation

In order to obtain optimal results with the gradient-magnitude method, several pre-processing steps should be performed on the data. Ranging from absolutely necessary to desired but not critical, these pre-processing steps include: (i) bias field correction, (ii) brain extraction, (iii) cerebellum removal and (iv) removal of brain stem structures. Successful bias field correction is critical to performance since otherwise intensity values for different tissue types start to mix in 2D histogram space. Brain extraction should be performed to remove irrelevant voxels from the 2D histogram representation. Removal of cerebellar and brain stem structures is recommended since it further improves conformity to ideal 2D histogram shapes (Fig 1D). Bias field correction and brain extraction can be performed using automatic algorithms [62, 63]. Removal of cerebellum and sub-cortical structures might require the manual creation of masks. We note, however, that generation of these masks is only desired, not strictly necessary. Furthermore, generation of these masks is often a desirable processing step for many automatic tissue class segmentation algorithm, since it improves their performance.

### 3.2 Data requirements

Suitability of the intensity-gradient magnitude histogram for separating brain from non-brain tissue will depend on the resolution and CNR of the input data. We expect a lower limit of resolution around 1 mm. At lower resolutions, the intensity-gradient magnitude method will yield unsatisfactory results due to partial voluming between the thin structures we are aiming to correct and surrounding CSF or tissues. We do not expect an upper resolution limit for the input data. Although, initially, values in the gradient magnitude image will no longer be high in all vessel and dura mater voxels, very high-resolution images can still be accommodated by choosing the appropriate level of smoothness on the gradient magnitude image. In S1 Fig, we demonstrate that by setting the appropriate smoothness level of Deriche filter [46], gradient magnitude images for very high resolution data (0.25 mm isotropic) [64, 65] can be approximated to those observed for data at lower resolution (0.7 mm isotropic).

We furthermore expect our methods to be impacted by the CNR of the input data. S2 Fig, S3 Fig and S4 Fig show that with added Gaussian noise (i.e. decreasing CNR) the desired circular and arc-like shapes in the 2D histogram (Fig 1C) become less apparent. At very high noise levels separating brain from non-brain tissue in the 2D histogram space becomes challenging (see e.g. S2 Fig). While the in-depth evaluation of additional processing tools is beyond the scope of the present article, we note that if the input data are very noisy, smoothing can be applied. In particular, non-linear anisotropic diffusion based smoothing [66, 67] results in the data regaining the desired 2D histogram shapes (see S5 Fig).

The parameter space of the input data is thus constrained by resolution and CNR. Apart from these restrictions, our methods are suitable for any 3D image and work irrespective of the field-of-view of the acquisition (partial coverage is possible) and membership to a particular species (bottle-nose dolphin brain is also possible; for examples see S6 Fig).

## 4 Validation methods

### 4.1 Validation data set overview

In order to validate the methods proposed above, we created two validation data sets based on the acquisition of high-resolution 7 T data of nine subjects and corresponding manually-guided expert segmentations of GM. In particular, we created two validation sets based of on two of the most common acquisition sequences. For five subjects, we collected MPRAGE T1w, PDw, and T2* data (we refer to this data set as the MPRAGE data set below). For four different subjects, we collected MP2RAGE data, to obtain unbiased (uni) images, and multi-echo 3D GRE data, to obtain T2* maps (we refer to this data set as the MP2RAGE data set below). Both data sets can be downloaded from [49].

#### 4.1.1 Ethics statement

The experimental procedures were approved by the ethics committee of the Faculty for Psychology and Neuroscience (MPRAGE data set) or the Medical Ethical Committee at the Faculty of Health, Medicine and Life Sciences (MP2RAGE data set) at Maastricht University, and were performed in accordance with the approved guidelines and the Declaration of Helsinki. Written informed consent was obtained for every participant before conducting the experiments.

#### 4.1.2 MRI acquisition parameters

All images were acquired on a Siemens 7 T whole body scanner (Siemens Medical Solutions, Erlangen, Germany) using a head RF coil (Nova Medical, Wilmington, MA, USA; single transmit, 32 receive channels). In all acquisitions, we used dielectric pads [68].

For *n*=5 subjects (age range 24-30, 2 females, no medical condition), the MPRAGE data set consisted of: a T1w image using a 3D MPRAGE sequence (repetition time [TR] = 3100 ms; time to inversion [TI] = 1500 ms [adiabatic non-selective inversion pulse]; time echo [TE] = 2.42 ms; flip angle = 5°; generalized auto-calibrating partially parallel acquisitions [GRAPPA] = 3 [69]; field of view [FOV] = 224 × 224 mm2; matrix size = 320 × 320; 256 slices; 0.7 mm isotropic voxels; pixel bandwidth = 182 Hz/pixel; first phase encode direction anterior to posterior; second phase encode direction left to right), a PDw image (0.7 mm isotropic) with the same MPRAGE as for the T1w image but without the inversion pulse (TR = 1380 ms; TE = 2.42 ms; flip angle = 5°; GRAPPA = 3; FOV = 224 × 224 mm; matrix size = 320 × 320; 256 slices; 0.7 mm iso. voxels; pixel bandwidth = 182 Hz/pixel; first phase encode direction anterior to posterior; second phase encode direction left to right), and a T2*w anatomical image using a modified MPRAGE sequence that allows freely setting the TE (TR = 4910 ms; TE = 16 ms; flip angle = 5°; GRAPPA= 3; FOV = 224 × 224 mm; matrix size = 320 × 320; 256 slices; 0.7 mm iso. voxels; pixel bandwidth = 473 Hz/pixel; first phase encode direction anterior to posterior; second phase encode direction left to right).

For *n*=4 subjects (age range 24-58, 2 females, no medical condition) the MP2RAGE data set consisted of: 3D MP2RAGE data (TR = 5000 ms; TI1/TI2 = 900/2750 ms; TE = 2.46 ms; FA1/FA2 = 5°/3°; FOV = 224×224 mm2; matrix size = 320 × 320; slices = 240; 0.7 mm iso voxels) [26]. For the same subjects, T2*w images were obtained with a multi-echo 3D GRE sequence (TR = 33 ms; TE1/TE2/TE3/TE4 = 2.53/7.03/12.55/20.35 ms; FA1 = 11°; FOV = 224 × 159 mm2; matrix = 320 × 227; slices = 208; 0.7 mm iso voxels). More details on the MP2RAGE data acquisition and the T2* estimation can be found in [70].

#### 4.1.3 Manually-guided expert segmentations

For every subject, we established “ground truth” GM classifications via manually-guided expert segmentations. All segmentations were created manually by the same expert (OFG), using ITK-SNAP [71] and a graphics tablet (Intuos Art; Wacom Co. Ltd; Kazo, Saitama, Japan). Segmentations were only established for cortical GM, since cerebellar and sub-cortical structures were later removed in a pre-processing step. To establish the segmentation, the expert used T1w images for the MPRAGE and uni images for the MP2RAGE data set. To avoid resulting tissue type classification to be ragged, the expert followed a particular processing sequence. The brain was first traversed in a single direction (e.g. sagittally) and the ground truth was established slice-by-slice. Subsequently, the brain was traversed in the two other directions (e.g. axially, then coronally). This sequence was repeated several times across several regions until the GM segmentation of the whole brain was considered of good quality. To further ensure the quality of the resulting segmentation, they were inspected for mistakes by two additional experts (MS and FDM).

### 4.2 Software implementation

We implemented the creation of transfer function based on 2D histograms in an open source Python package called Segmentator [48], which is built upon several other scientific packages such as Numpy [72], Scipy [73], Matplotlib [74] and Nibabel [75]. Segmentator allows for selection of data points in a 2D histogram (for example gradient magnitude over intensity) and concurrent visualization of selected brain voxels on a 2D slice. Data points can be selected using a circular sector widget with variable reflex angle and radius. Alternatively, data selection can be performed using the normalized graph cut (n-cut) method (i.e. spectral clustering) as described above. The n-cut algorithm from Scikit-image [76] was modified to export an augmented output which provides step-wise access to independent branches of the decision tree and employed in Segmentator (the modification is available at: https://github.com/ofgulban/scikit-image/tree/ncut-rag-options).

The package provides several options to calculate the gradient magnitude image. All the 2D histogram analyses described in this paper were based on gradient magnitude images that were computed as the Euclidean norm of the first spatial derivative estimated using a 3 × 3 × 3 Scharr kernel [77, 78]. Subsequently, transfer functions were specified using the normalized graph cut algorithm and user intervention for the selection of the non-brain tissue transfer functions. Processing data for a single subject took about 10 minutes on average. The Segmentator package is openly and freely accessible at https://github.com/ofgulban/segmentator. All the operations of the CoDa analysis described above have been implemented as a separate open source Python package [79] freely accessible at https://github.com/ofgulban/compoda. This package uses Numpy [72] and Scipy [73].

### 4.3 Segmentation procedure

For both validation data sets, we followed similar procedures, with modifications where necessary to accommodate for differences in the sequences’ output. Our goal was to obtain initial GM segmentations from existing, fully-automated segmentation algorithms and to quantify the improvement in segmentation accuracy that can be obtained when using the methods described here as post-processing steps. To establish the initial GM segmentations we used FSL FAST [80] and the SPM 12 “unified segmentation” algorithm [63] for the MPRAGE data set and FSL FAST and CBS tools [32] for the MP2RAGE data set. SPM and CBS tools have been developed and benchmarked on MPRAGE and MP2RAGE images respectively. FSL FAST is suited to process either type, so we used it for both data sets. We then quantified the impact of the following additional post-processing steps: (i) using uni-modal input and transfer functions based on 2D histogram representations of intensity and gradient magnitude (see Section 1) or (ii) using multi-modal input and the compositional data analysis framework (see Section 2). These two procedures will be referred to below as the gradient magnitude (GraMag) and the compositional data analysis (CoDa) method, respectively. Both methods resulted in masks that could be used to further refine the initial GM segmentation, e.g. by removing blood vessels and dura mater that were falsely labeled as GM initially. In total, we thus used 2 (MPRAGE and MP2RAGE data set) × 2 (GraMag and CoDa) = 4 analysis procedures. All four procedures are summarized in flow chart diagrams (S8 Fig, S9 Fig, S10 Fig, S11 Fig). Furthermore, in an effort to make our analyses fully reproducible, we made the Python and bash scripts used for pipeline processing openly available at [50].

For the MPRAGE data set, we first computed ratio images (T1w divided by PDw) [25] to reduce inhomogeneities. Ratio images were input to either FSL FAST or SPM 12. FSL FAST was used with default values. The FAST algorithm requires an initial brain extraction procedure that we performed using FSL BET [62]. Additionally, we masked the images to exclude: the corpus callosum, the basal ganglia, the hippocampus, the entire brain stem and the cerebellum. Below we refer to this mask as “NoSub mask”. The NoSub mask was created manually for every subject. In SPM 12 we used default settings with one exception. We set the number of Gaussians to be modeled to 3 for GM and 2 for WM (default values are 1 and 1). As part of their standard segmentation routine, both FSL FAST and SPM 12 perform initial inhomogeneity correction. We output and inspected the bias corrected images to ensure that the algorithms had converged on plausible solutions. We specified for the FSL FAST algorithm to output hard segmentation labels. Since SPM 12 outputs probabilities for six tissue classes, we transformed this soft output to hard segmentation labels by assigning each voxel to the tissue class with the highest posterior probability. Since the SPM segmentation algorithm works best with unmasked images, we applied the NoSub mask only to the resulting SPM GM segmentations, not to the input data. The resulting GM segmentations from FSL and SPM were saved for later evaluation.

For the GraMag method (S8 Fig) we proceeded with bias-corrected ratio images from either SPM or FSL. Since the GraMag method works best with brain extracted images, we combined SPM’s WM and GM segmentation outcomes to form a brain mask and performed brain extraction of the ratio images from SPM. After brain extraction, we also excluded cerebellum and brain stem tissue using the NoSub mask. FSL’s bias-corrected ratio images images did not require masking as the brain extraction (and cerebellum removal) was already performed before segmentation. We then used the 2D histogram representation of intensity and gradient magnitude together with the hierarchical exploration of normalized graph cut decision trees (as described in Section 1) to create transfer functions. Exploration of decision trees was limited to an 8-level hierarchy. The criterion for splitting and merging clusters was subjective: a rater (MS) aimed to obtain shapes that resembled the ideal template shapes (Fig 1D) as closely as possible, given the 2D histogram representation and concurrent visualization of selected voxels. S1 Video demonstrates that selection of voxels was well constrained by clearly-outlined shapes in the 2D histogram representation and commonly required to move down the decision tree hierarchy by only 2-3 levels. Exploration of the decision tree took about 30 - 60 seconds per subject. Generation of normalized graph cut decision trees, which was done previous to exploration by a rater, took about 5 minutes on a workstation computer (RAM: 32 GB, 12 cores (6 virtual); CPU: 2.146 GHz; operating system: Debian 8). The transfer function resulting from this procedure was used to separate brain from non-brain tissue voxels. Non-brain tissue voxels were removed from GM if they were included in the initial FSL and SPM segmentations.

For the CoDa method (S9 Fig) we followed a similar procedure, except that we started from three separate images - the bias-corrected T1w, PDw and T2*w images. Again, these images were brain extracted and cerebellum and brain stem tissue were removed using the NoSub mask. These images were transformed into barycentric coordinates, using the closure operator (as outlined in in Section 2). In this case, there were three barycentric coordinates per voxel constrained to a 2-simplex vector space structure. The triplets of barycentric coordinates were mapped to 2D real-space using the *ilr* transformation. We could therefore proceed with the 2D histogram representation using the first and the second real-space coordinates of the compositions and the hierarchical exploration of normalized graph cut decision trees in this 2D space to separate brain from non-brain tissue voxels. Non-brain tissue voxels were again removed from GM if included in the initial segmentations (of SPM and FSL).

For the MP2RAGE data set, the T1 map, T1w (uni) and second inversion image from the MP2RAGE sequence were input to CBS tools [32]. Only the brain-extracted [62] uni image was input to FSL FAST, since this resulted in higher performance than inputting all three images. Both FSL FAST and CBS tools were run with default settings. Note that the default settings for CBS tools include removal of non brain tissue by estimating dura mater and CSF partial voluming. The resulting GM segmentations from FSL and CBS were saved for later evaluation. For the GraMag method (S10 Fig), we proceeded with the FSL FAST bias-corrected, brain-extracted and NoSub masked uni image and proceeded as for the MPRAGE data set to obtain a secondary brain mask. For the CoDa method (S11 Fig), we used FSL FAST bias-corrected, brain-extracted and NoSub masked uni, second inversion and T2* images but otherwise proceeded as for the MPRAGE data set.

We observed susceptibility artifacts in some regions of the brain (mostly inferior frontal lobe) in the T2*w images. These artifacts make the affected regions noisy and reduce the effectiveness of using T2*w images in the CoDa method. To quantify the effect of these artifacts, we created masks for the artifact-affected regions and ran all our analyses both with and without the artifact regions included. Results shown in the paper were obtained with the affected regions excluded. Results with the affected regions included are shown in the Supplementary Materials.

### 4.4 Quantification

The segmentation procedures resulted in three different GM segmentations for each data set and initial segmentation algorithm (SPM or CBS and FSL FAST): (i) an initial segmentation without any further changes, (ii) after correction using the GraMag method and (iii) after correction using the CoDa method. To compare segmentation quality among these three outcomes, we calculated the Dice coefficient (DICE) and the Average Hausdorff Distance (AVHD) using the openly available EvaluateSegmentation Tool (2016; VISCERAL, http://www.visceral.eu).

The DICE is an overlap-based metric and it is the most popular choices for validating volume segmentations [81]. We included it here as a familiar reference for the reader. However, overlap-based metrics like the DICE are not recommended for validating segmentation boundaries against the ground truth, as is our aim here, since they are relatively insensitive to boundary errors. In contrast, the AVHD is a distance-based metric and is sensitive to boundary errors [81]. We therefore consider the AVHD to be a more suitable metric for our purposes and we based our conclusions on the comparisons made with the AVHD.

Given that the AVHD quantifies the similarity of two boundaries, we first extracted WM-GM and GM-CSF boundaries from the ground truth segmentations and the six different GM segmentations before calculating the AVHD. Here, an AVHD of 0 indicates a perfect match between the segmentation and ground truth boundaries, while values >0 indicate a mismatch. In this case, the value represents the average number of voxels by which the two boundaries deviate from one another. For example, an AVHD of 1 indicates that the segmentation boundary, on average, deviates of one voxel from the ground truth.

### 5 Validation results

Visual inspection revealed that applying the GraMag method to the MPRAGE data set excluded most of the vessels and dura mater voxels and resulted in a more plausible GM matter definition. The CoDa method equally removed most of the vessels and dura mater voxels. Additionally, the CoDa method excluded structures like the sagittal sinus from the GM definition (see Fig 6 and Fig 7).

**Fig 6.**
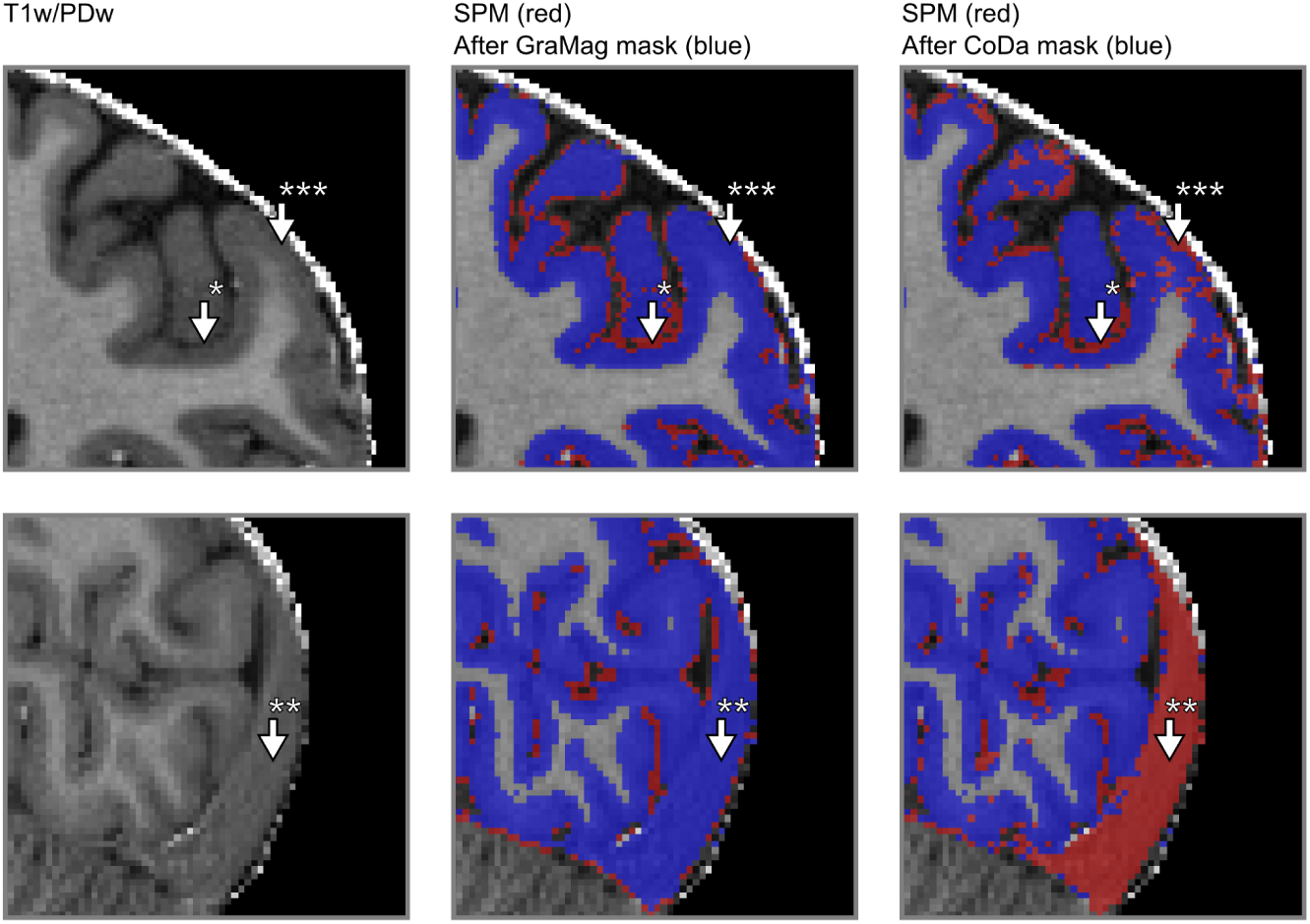
Comparison of GM segmentation results for MPRAGE data. GM segmentation results are shown for one representative subject on a transverse (upper row) and a sagittal slice (lower row) of the brain before and after applying the GraMag and CoDa methods. The original image that is input to the segmentation is shown on the left. The original GM segmentation obtained from SPM 12 is shown in red (middle and right column). GM segmentations after additional polishing with brain mask obtained with either the GraMag (middle column) or the CoDa method (right column) are overlaid in blue. Additional masking removes blood vessels, CSF (arrow *) and most of dura mater (arrow †) voxels from the SPM GM definition. Because of its unique compositional properties, connective tissue from the sagittal sinus can be captured and excluded using the CoDa method (arrow **). An area badly affected by the CoDa mask is also indicated with arrow ***.

**Fig 7.**
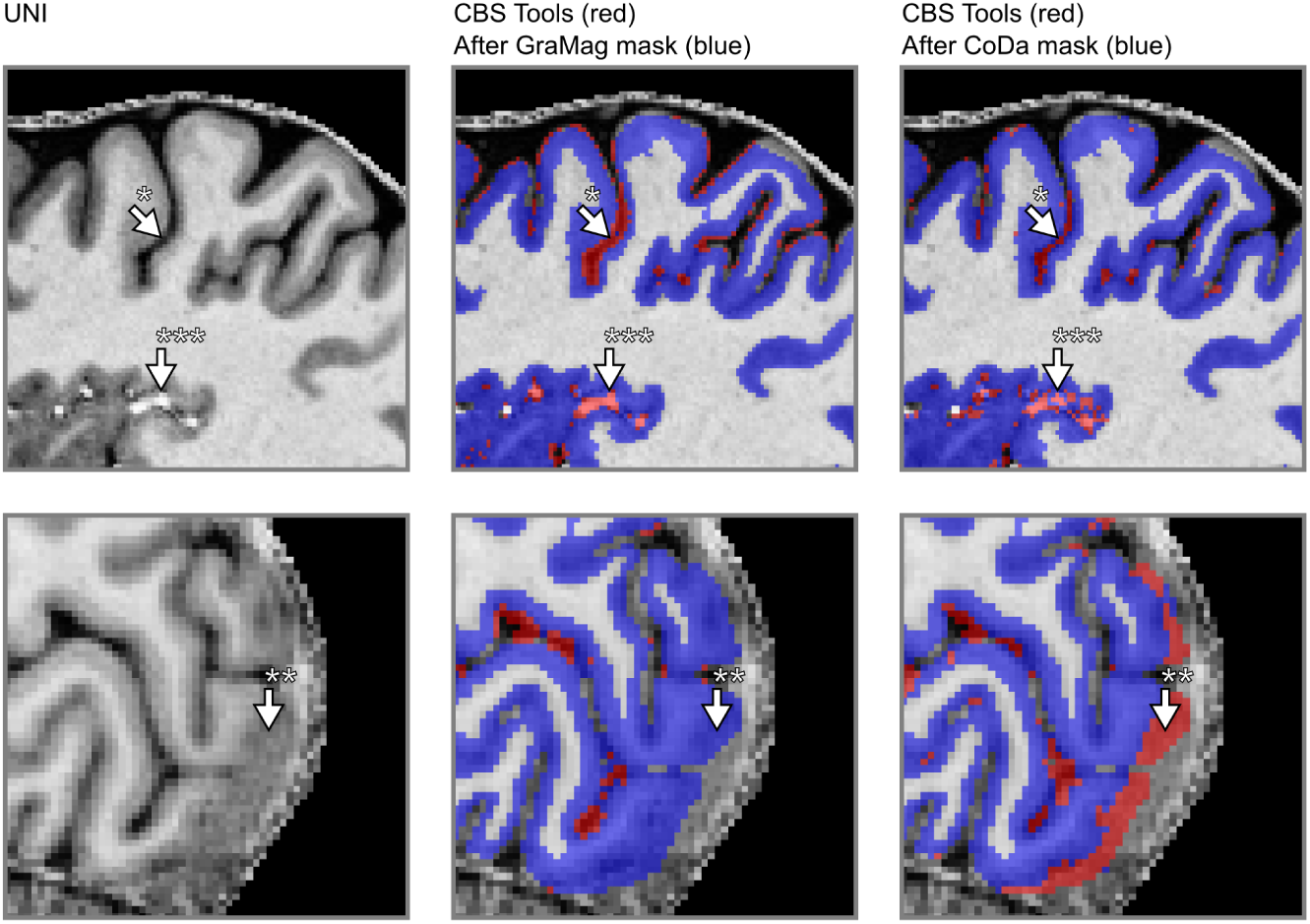
Comparison of GM segmentation results for MP2RAGE data. Same conventions as in Fig 6) but with initial segmentation results obtained with CBS tools instead of SPM 12.

Table 2 compares segmentation performance before and after applying GraMag and CoDa methods to the initial GM segmentations of the MPRAGE data set. The GraMag method led to an improvement of GM segmentations in all subjects, independently of whether the initial segmentation was done by SPM 12 or FSL FAST. On average, the AVHD decreased from 0.733 ±0.087 (mean ±standard deviation across subjects) to 0.571±0.051 for SPM 12 and from 0.584 ±0.109 to 0.558 ±0.089 for FSL FAST. The GraMag method did not affect the DICE coefficient. On average, it changed very little from 0.861 ±0.020 to 0.862 ±0.016 for SPM 12 and from 0.878 ±0.027 to 0.872 ±0.089 for FSL FAST. The CoDa method equally yielded improved segmentation performance. Compared to the initial segmentation, the AVHD decreased in all subjects and, on average, to 0.569 ±0.054 for SPM 12 and to 0.504 ±0.033 for FSL FAST. We did not observe a clear change in the DICE coefficient. For SPM 12 segmentations we observed 0.869 ±0.021 and for FSL FAST segmentations 0.872 ±0.013. All these results were obtained after exclusion of areas affected by artifacts in the T2s image. For results obtained without the artifact masks, please see S1 Table.

**Table 2.** Segmentation performance scores MPRAGE data set. The table shows the DICE (larger is better) and AVHD (less is better) for the initial SPM 12 and FSL FAST GM segmentations as well as after additional polishing, using either the gradient magnitude or the compositional data method.

|  |  | SPM |  | FAST |  |
| --- | --- | --- | --- | --- | --- |
|  |  | DICE<sup>a</sup> | AVHD<sup>a</sup> | DICE<sup>a</sup> | AVHD<sup>a</sup> |
| S02 | Init | 0.8606 | 0.7325 | 0.8869 | 0.5661 |
|  | Init + GraMag | 0.8613 | 0.5811 | 0.8876 | 0.5162 |
|  | Init + CoDa | 0.8689 | 0.5693 | 0.8719 | 0.5017 |
| S03 | Init | 0.8398 | 0.8585 | 0.8782 | 0.5835 |
|  | Init + GraMag | 0.8679 | 0.6031 | 0.8722 | 0.5575 |
|  | Init + CoDa | 0.8625 | 0.5986 | 0.8723 | 0.5039 |
| S05 | Init | 0.8681 | 0.6350 | 0.8858 | 0.5545 |
|  | Init + GraMag | 0.8595 | 0.5618 | 0.8770 | 0.5359 |
|  | Init + CoDa | 0.8796 | 0.4950 | 0.8960 | 0.4601 |
| S06 | Init | 0.8428 | 0.7778 | 0.8633 | 0.6462 |
|  | Init + GraMag | 0.8624 | 0.5711 | 0.8665 | 0.5848 |
|  | Init + CoDa | 0.8592 | 0.5880 | 0.8775 | 0.5074 |
| S07 | Init | 0.8945 | 0.6205 | 0.8583 | 0.6925 |
|  | Init + GraMag | 0.8983 | 0.5281 | 0.8623 | 0.6487 |
|  | Init + CoDa | 0.8692 | 0.5541 | 0.8248 | 0.6689 |

DICE, DICE Coefficient; AVHD, Average Hausdorff Distance; Init, initial segmentation; GraMag, gradient magnitude method; CoDa, compositional data method.
<sup>a</sup> After masking of areas affected by artifact in the T2s image.

Table 3 compares segmentation performances before and after applying the GraMag and CoDa methods to the initial GM segmentations of the MP2RAGE data set. The GraMag method decreased AVHD for all but one subject and, on average, from 0.508 ±0.088 to 0.444 ±0.083 for CBS tools and from 0.990 ±0.062 to 0.775 ±0.088 for FSL FAST. It decreased the DICE coefficient, on average, from 0.882 ±to 0.875 ±for CBS tools and from 0.839 ±to 0.818 ±for FSL FAST. The CoDa method decreased the AVDH in every subject and, on average, to 0.447 ±0.082 for CBS tools and to 0.641 ±0.069 for FSL FAST. It also increased the DICE coefficient to 0.856 ±0.030 for FSL FAST and decreased it to 0.880 ±0.024 for CBS tools. For results for the MP2RAGE data obtained without the artifact masks, please see S2 Table.

**Table 3.** Segmentation performance scores MP2RAGE data set. The table shows the DICE (larger is better) and AVHD (less is better) for the initial CBS tools and FSL FAST GM segmentations as well as after additional masking, using either the gradient magnitude or the compositional data method.

|  |  | CBS |  | FAST |  |
| --- | --- | --- | --- | --- | --- |
|  |  | DICE<sup>a</sup> | AVHD<sup>a</sup> | DICE<sup>a</sup> | AVHD<sup>a</sup> |
| S001 | Init | 0.8943 | 0.3711 | 0.8403 | 1.0523 |
|  | Init + GraMag | 0.9141 | 0.4013 | 0.8248 | 0.8241 |
|  | Init + CoDa | 0.9185 | 0.3249 | 0.8795 | 0.5757 |
| S013 | Init | 0.8672 | 0.5641 | 0.7859 | 1.0272 |
|  | Init + GraMag | 0.8629 | 0.4921 | 0.7653 | 0.8835 |
|  | Init + CoDa | 0.8648 | 0.4835 | 0.8137 | 0.7225 |
| S014 | Init | 0.8695 | 0.5897 | 0.8376 | 0.9203 |
|  | Init + GraMag | 0.8606 | 0.4859 | 0.8136 | 0.7260 |
|  | Init + CoDa | 0.8710 | 0.4921 | 0.8486 | 0.6789 |
| S019 | Init | 0.9077 | 0.4517 | 0.8529 | 0.9529 |
|  | Init + GraMag | 0.8865 | 0.3301 | 0.8218 | 0.7005 |
|  | Init + CoDa | 0.8888 | 0.4096 | 0.8642 | 0.6024 |

DICE, DICE Coefficient; AVHD, Average Hausdorff Distance; Init, initial segmentation; GraMag, gradient magnitude method; CoDa, compositional data method.
<sup>a</sup> After masking of areas affected by artifact in the T2s image.

### 6 Discussion

Functional and anatomical MRI studies at the mesoscale (< 1 mm isotropic) require accurate and precise definitions of the GM ribbon. Creating such definitions is currently a challenging task since sub-millimeter UHF data bring non-brain structures like blood vessels and dura mater into sharper focus. As a result, segmentation algorithms that have been benchmarked at lower resolution data might falsely label part of these structures as GM. Here we presented two methods (GraMag and CoDa) to correct many such mislabeled non-brain voxels efficiently and semi-automatically. The two methods are based on theoretical expectations of how 3D brain data is to be represented in 2D histograms. These expectations imply that brain and non-brain tissue should become separable in 2D histogram representations that are either based on gradient magnitude and intensity or on compositional dimensions. We validated these expectations by implementing the suggested methods in an openly available software package and by quantifying their added benefit using a new high-resolution validation data set. We found that, in general, our suggested methods offered an improvement compared to initial GM segmentations. However, we found some differences in the degree of improvement with respect to (i) the two presented methods, the (ii) type of data and (iii) the algorithm used for initial segmentation.

We will discuss these three influences in turn. First, the two methods differ in their prerequisites and their segmentation improvement. The GraMag method only requires uni-modal input such as T1w/PDw or MP2RAGE uni images, while the CoDa method requires multi-modal input of images with different contrast weightings. This makes the GraMag method the method of choice when only a single input image is available. In accordance with our theoretical expectation, the GraMag method identified and removed blood vessels and dura mater tissue. If multi-dimensional input is available, even bigger improvements might be obtained with the CoDa method. Notably, in contrast to the GraMag method, the CoDa method can additionally capture and remove connected tissue of the sagittal sinus. This tissue is usually falsely labeled as GM because of similar intensity values and spatial proximity. It then requires tedious manual removal. How well the CoDa method performs, however, critically depends on the quality of all the input images and the specific combination of contrasts. Performance can be affected by low quality on a single input image, as was the case here with T2* images due to susceptibility artifacts. Furthermore, performance will depend on the specific choice of contrasts and whether these contrasts maximize the compositional difference between brain and non-brain tissue.

Second, we found that the improvements were slightly larger and more consistent across subjects for the MPRAGE than for the MP2RAGE data set. This might be explained by the fact that the MPRAGE data conformed more to our theoretical expectations than the MP2RAGE data set. Especially, we found GM values in the MP2RAGE uni image to be less focused on one particular area of the 2D histogram (S12 Fig) than the MPRAGE division image. This might result from differences in myelination level across cortical areas and depth [54, 82, 83], which the MP2RAGE uni image might pick up more than MPRAGE division image [84].

Third, we observed that the performance of the initial segmentation algorithm had an influence on how much we could further improve the GM segmentation. If performance of the initial segmentation algorithm was already relatively high, the improvement obtained with our methods tended to be smaller. Differences in initial segmentation performance might be explained by whether the algorithm has been benchmarked on this particular type of data. We assume FSL FAST and CBS tools to have been benchmarked on MPRAGE and MP2RAGE data respectively, which would explain their relative high performance for these data types.

Importantly, our goal here was to aid already existing segmentation pipelines to deal with UHF sub-millimeter resolution data, not to replace those pipelines. Instead, the methods presented here should be considered as an alternative to a large amount of manual slice-by-slice polishing of segmentations and thus as a time-saver. Manually correcting segmentation labels is very time-consuming and can quickly become unreliable. In contrast, our methods greatly reduce the time required for manual polishing because they offer an efficient 2D summary and are more reliable because they are semi-automatic. Although the methods presented here do not entirely eliminate the need for manual corrections, we estimated that for a whole brain cortical ribbon segmentation they do save on average 7.5 hours of manual work (for more details on this estimation see S1 Appendix).

Moreover, we introduced the compositional data analysis framework to the neuroimaging community. Here, we used this framework to combine MRI acquisitions with three different image contrasts in order to derive improved tissue type segmentations. While the compositional analysis framework scales to any dimension and thus any number of MRI images, the current implementation relies on representation of data in a 2D histogram obtained through the ilr transformation of 3D barycentric coordinate data. With more than three images a reduction of dimensions in the barycentric space or in the real space after ilr transformation would be necessary to apply the current tools (e.g. [40]).

MRI can provide a multitude of informative images that weight tissue properties to generate the image contrast. The compositional data framework is ideally suited for the analysis and visualization of multiple images as it provides a principled way to combine any number of images. In addition, analyzing multiple MRI contrast images in the compositional data framework avoids spatial scale dependence, i.e. dependence on the image resolution and smoothness of the image. As a result, the compositional properties of vessel voxels even at very high resolutions will remain the same or very similar, no matter whether the voxel is at the center or at the border of the vessel. This is similar to analyzing chemical compositions of materials, which are independent of spatial metrics.

An envisioned future application of the compositional framework to MRI data is to use it to single out targeted cortical or subcortical structures based on their compositional properties. For an example of identifying subcortical structures see S7 Fig. For discussion of the broader implications of the application of compositional data analysis to images in general please see [85].

Our theoretical expectations implied that the methods presented here require high-resolution data (<1 mm). This requirement was unfortunately not met by most available segmentation validation data sets. Simulated phantom (“BrainWeb”) data [86] are available at 1 mm and thus fell short of the resolution required for our purposes. Although an updated data set (“updated BrainWeb”, [87, 88]) is available at higher resolution, the simulations in this data set were based on initial 3T MRI acquisitions. As a consequence, the updated BrainWeb data revealed considerably less bright vessel and dura mater voxels than 7 T data usually does and was not suitable to validate our methods.

These considerations led us to create our own high-resolution segmentation validation data sets for which we established the “ground truth” via manually-guided expert segmentation. While expert segmentations have well-known drawbacks [33, 89], they also have important advantages to alternative methods of establishing the ground truth, such as simulated phantom data. In particular, creating a validation data set based on empirical data and expert segmentations allowed us to benchmark our methods under conditions where image intensities fell into the expected range. Being aware of the problems with expert segmentations, we alleviate concerns about the quality of our expert judgment and consequently the validity of the results presented here by taking the following measures. First, the final ground truth segmentations were inspected by two additional experts. Second, we make the data sets and corresponding ground truth segmentations as well as our processing scripts available. This will allow other researchers to come up with their own judgment of the quality of the ground truth segmentation and validation data. In case changes to the ground truth are suggested and implemented, quantification could be re-run using our openly-accessible work flow.

The 2D histogram method presented here is, in principle, capable of generating its own exhaustive tissue-type classifications, i.e. it does not necessarily depend on existing segmentation pipelines to derive GM and WM labels. While we expect the 2D histogram method to give no advantage over existing, fully-automated segmentation algorithms under standard conditions, the histogram method will compare well in cases where standard algorithms fail. Importantly, the 2D histogram method used here does not assume the data to conform to any atlas or template shape. Therefore, it is suitable also for acquisitions with only partial coverage (surface coils) or for specific populations like infant or even dolphin brains (see S6 Fig).

Using histogram-based methods would be more attractive if the process of specifying transfer functions was fully automatic. We note that there is no principled obstacle to doing this. Indeed, information-theoretic measures have been suggested [39] that would make the normalized graph-cut application fully automatic, given the specification of an appropriate stopping criterion. The transfer functions (i.e. the circles and arcs applied to our 2D histograms) that we observed for the different brain tissue types were stable across subjects and conformed to expected, ideal shapes. This would allow to define probabilistic templates in the histogram space and transform the methods proposed here to a fully automatic exhaustive tissue-type classifications.

We understand our methods as a secondary, more fine-grained brain extraction. When performing the initial brain extraction or tissue class segmentation, the user can often set parameters of the masking to be either more restrictive (at the risk of excluding brain tissue) or more liberal (at the risk of including a lot of non-brain tissue). We assume that, faced with this trade-off, users will usually lean to the liberal choice of parameters to avoid that relevant brain tissue is excluded. In such cases, we suggest our methods will prove useful. Our methods go beyond simply choosing more restrictive parameters because they focus on information that is relevant to excluding vessels, dura mater and connective tissue (Fig 1D).

Our comparisons were limited to segmentations obtained from FSL, SPM and CBS tools. While several MRI studies at the mesoscale have used alternative ways of establishing tissue class segmentations [90, 91], we decided to limit our comparison to openly available algorithms. Furthermore, the resolution of our validation data exceeded the recommended input range for FreeSurfer (1 mm to 0.75 mm isotropic).

As is to be expected, we found our methods to be be impacted by the CNR of the input data (S2 Fig - S4 Fig). In particular, additional noise caused the 2D histogram representation for both methods to conform less to expected template shapes. However, we note that for images that were acquired with currently very common imaging parameters at ultra high fields, we found our methods to offer clear benefits in GM segmentations. Furthermore, in case acquisitions are noisier than the ones tested here, additional processing steps like advanced smoothing [66, 67] might be applied to mitigate noise issues (see S5 Fig).

By making our validation data sets publicly available, we hope to inspire further algorithmic testing and development. There is currently a lack of validation data for the performance of tissue-type classification of MRI data acquired at ultra-high fields with sub-millimeter resolution. By publishing our data, our code and our work flow, we invite fellow scientists to benefit from our work but also to further contribute to it. The neuro-imaging community can use our data to test the performance of entirely new methods or modifications to existing segmentation algorithms. Contributions could be made in the form of additional high-resolution data, more ground truth segmentations and algorithmic improvement. Anticipating such algorithmic improvements, we envision a future where segmentation of volumetric images will become gradually less laborious despite increasing resolution and volume of the data.

## Acknowledgments

We would like to express our appreciation for the comments and suggestions of the reviewers which helped to improve the manuscript. We would also like to thank Thomas Emmerling for his guidance and advise on software implementation as well as Nikolaus Weiskopf, Rainer Goebel and Christophe Lenglet for valuable discussions of our early ideas. This work was financed by the Netherlands Organisation for Scientific Research (NWO). The authors O.F.G. and F.D.M. as well as data acquisition for the MPRAGE data set were supported by NWO VIDI grant 864-13-012. Author M.S. was supported by NWO research talent grant 406-14-108. Author I.M. was supported by NWO research talent grant 406-14-085. Author R.H. and acquisition of the MP2RAGE was funded by Technology Foundation STW (grant 12724).

## Supporting information

### S1 Appendix

#### Time benefit estimation

To estimate the time saved by substituting manual correction with Segmentator, we did the following calculation: Assuming that the objective of manually correcting mislabeled GM voxels is to reduce the number of false positives and to increase the number of true positives, we computed the number of false and true positives both before and after Segmentator intervention. We computed the difference in true positives and false positives before and after the application of segmentator and subtracted the true positive difference from the false positive difference. The resulting number is assumed to indicate the number of voxels which would have to be subtracted in the process of manually correcting all the voxels without using Segmentator. For the MRI data presented here, this number amounted to around 200000 voxels to be corrected (out of a total number of 1100000 voxels in the cortical ribbon). On average 35000 out of those 200000 voxels could be corrected using Segmentator. Assuming that a trained operator can manually correct one voxel per second, on average, this amounts to 7.5 hours of manual work that can be substituted with 10 minutes of Segmentator usage for the whole brain GM ribbon segmentation at 0.7 mm isotropic resolution. The time that can potentially be saved by using Segmentator will scale with the total number of GM voxels - it will be higher for high resolution acquisition (more voxels) and lower at low resolution (less voxels). The script used for our estimation is available at [50].

**S1 Fig.**
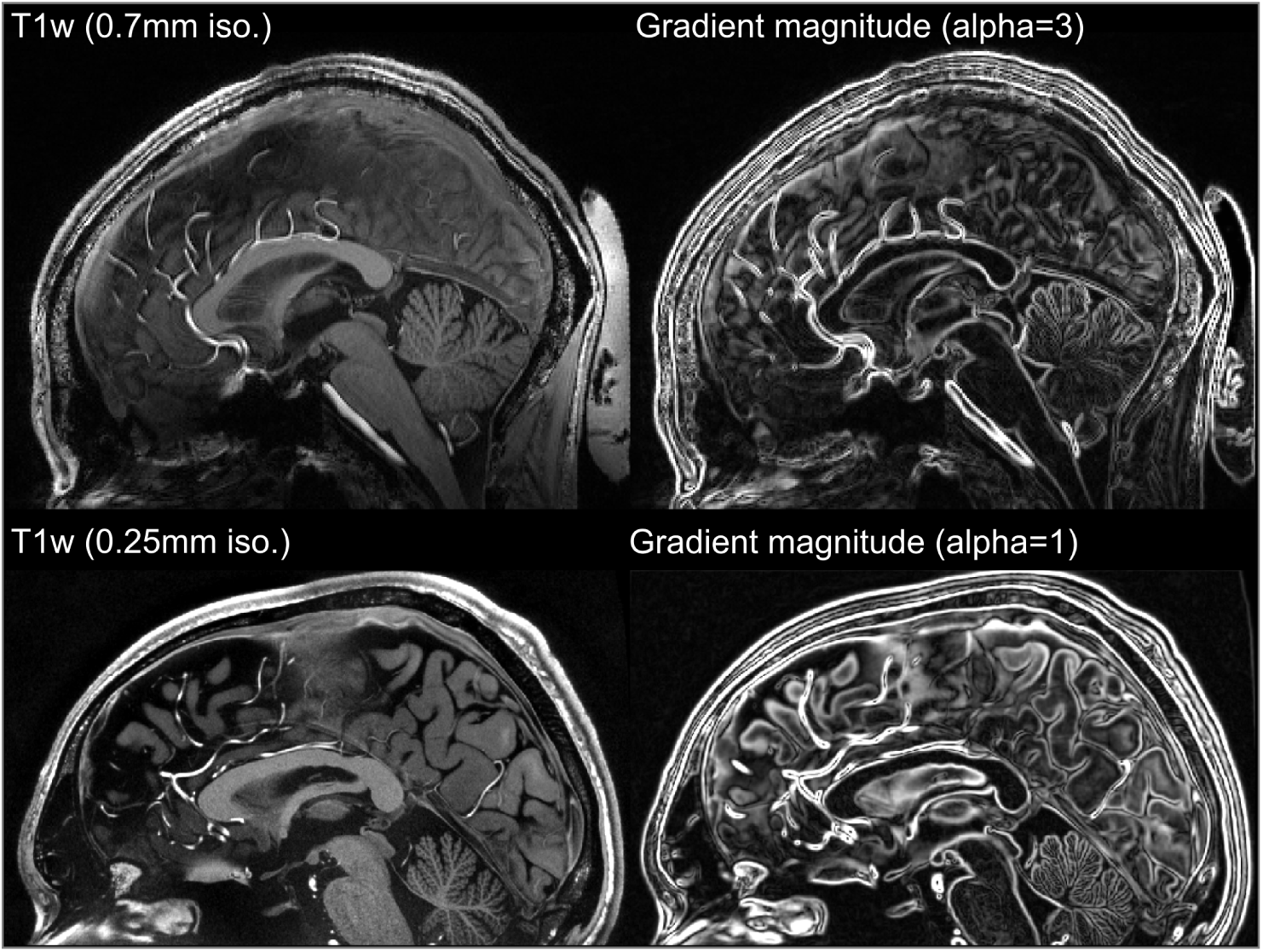
Appropriate kernel width approximates lower resolution. Intensity (left) and gradient magnitude (right) images are shown for T1w MRI data of a human brain that was either acquired at 0.7 mm isotropic (top) or at the 0.25 mm isotropic [64,65] (bottom). By choosing an appropriate kernel width for the very high resolution image (here alpha=1), the gradient magnitude image can be approximated to the lower resolution image, thus making it possible to use the gradient magnitude method also for very high resolutions. All images show a sagittal slice for an exemplary subject (sub-02).

**S1 Video. Using normalized graph cut decision trees for MRI data.** The exploration of normalized graph cut decision trees allows for finding a more restrictive brain mask that excludes dura mater and brain vessels in a quick and intuitive manner.

**S2 Fig.**
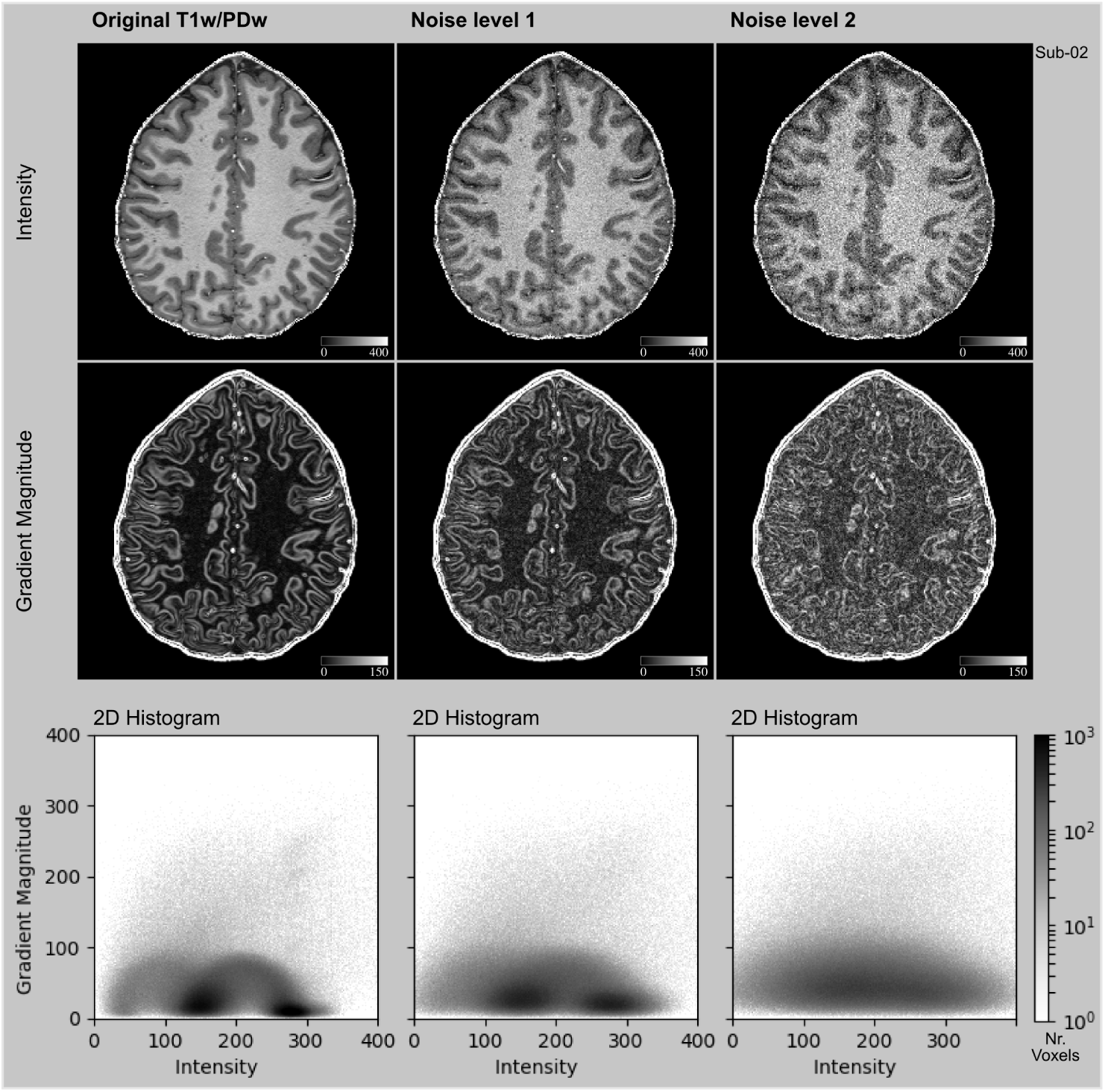
Impact of additional noise on GraMag method. Shown are intensity images (top row), gradient magnitude images (middle row) and 2D histograms for the GraMag method (bottom row) for a T1w-divided-PDw MRI ratio image without any additional noise (left) and after applying a moderate (middle) and high (right column) amount of additive Gaussian noise with two levels of constant standard deviation of the distribution. The moderate noise (*σ* = 25) is 16% and high noise (*σ* = 50) is 32% calculated relative to the mean cortical gray matter intensity. Noise causes structures in the 2D histogram that are initially well-defined to spread outward and, at very high noise levels, to lose shape. Images show a transverse slice for an exemplary subject (sub-02).

**S3 Fig.**
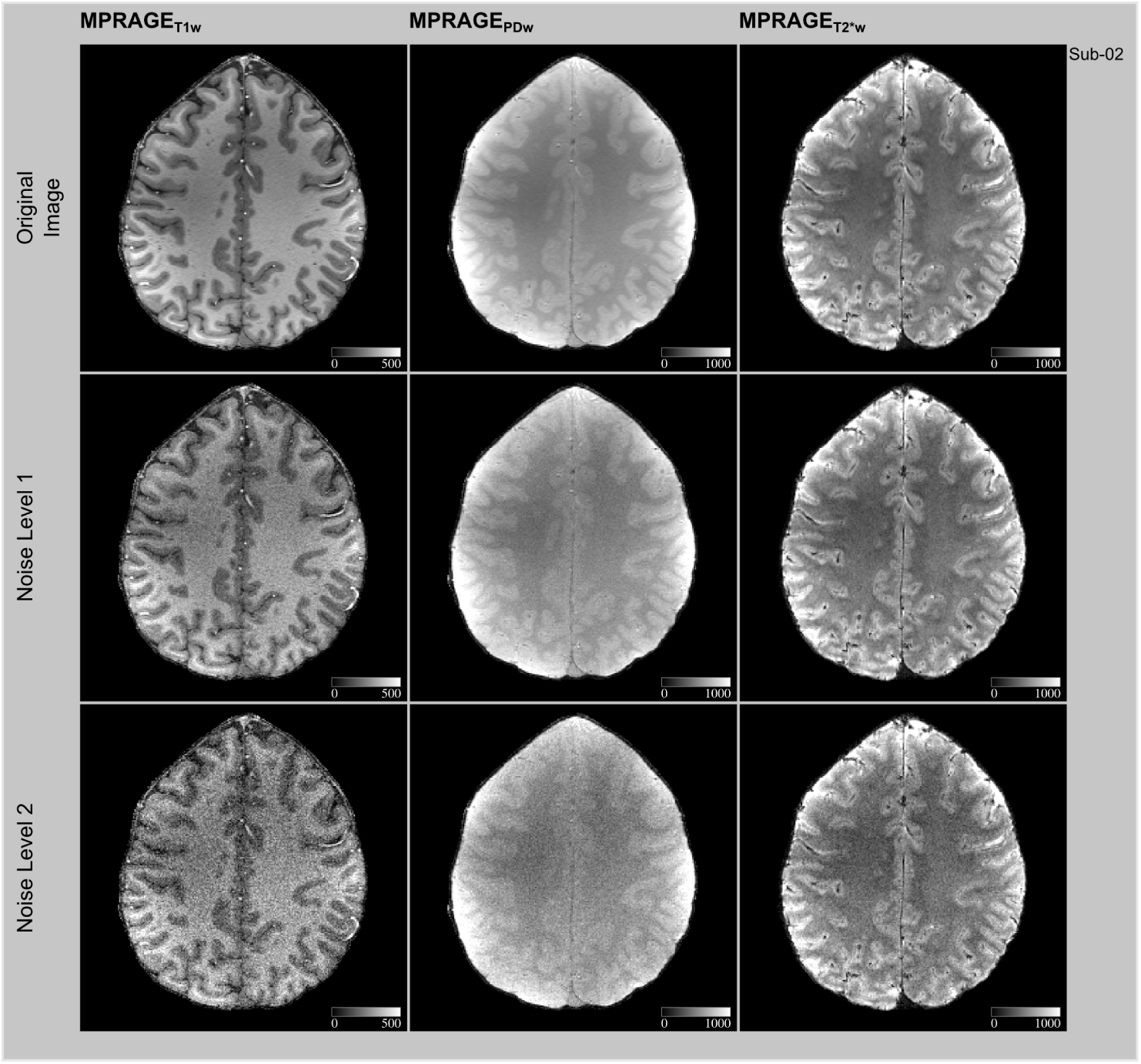
Impact of additional noise on CoDa method I. Shown are T1w (left), PDw (middle) and T2*w (right column) without any additional noise (top) and after applying a moderate (middle) and high (bottom row) amount of additive Gaussian noise with two levels of constant standard deviation of the distribution. Moderate noise (*σ* = 25) is 13% for T1w, 4% for PDw, 5% for T2*w calculated relative to the mean cortical gray matter intensity. High noise (*σ* = 50) is 27% for T1w, 9% for PDw, 10% for T2*w calculated relative to the mean cortical gray matter intensity. Images show a transverse slice for an exemplary subject (sub-02).

**S4 Fig.**
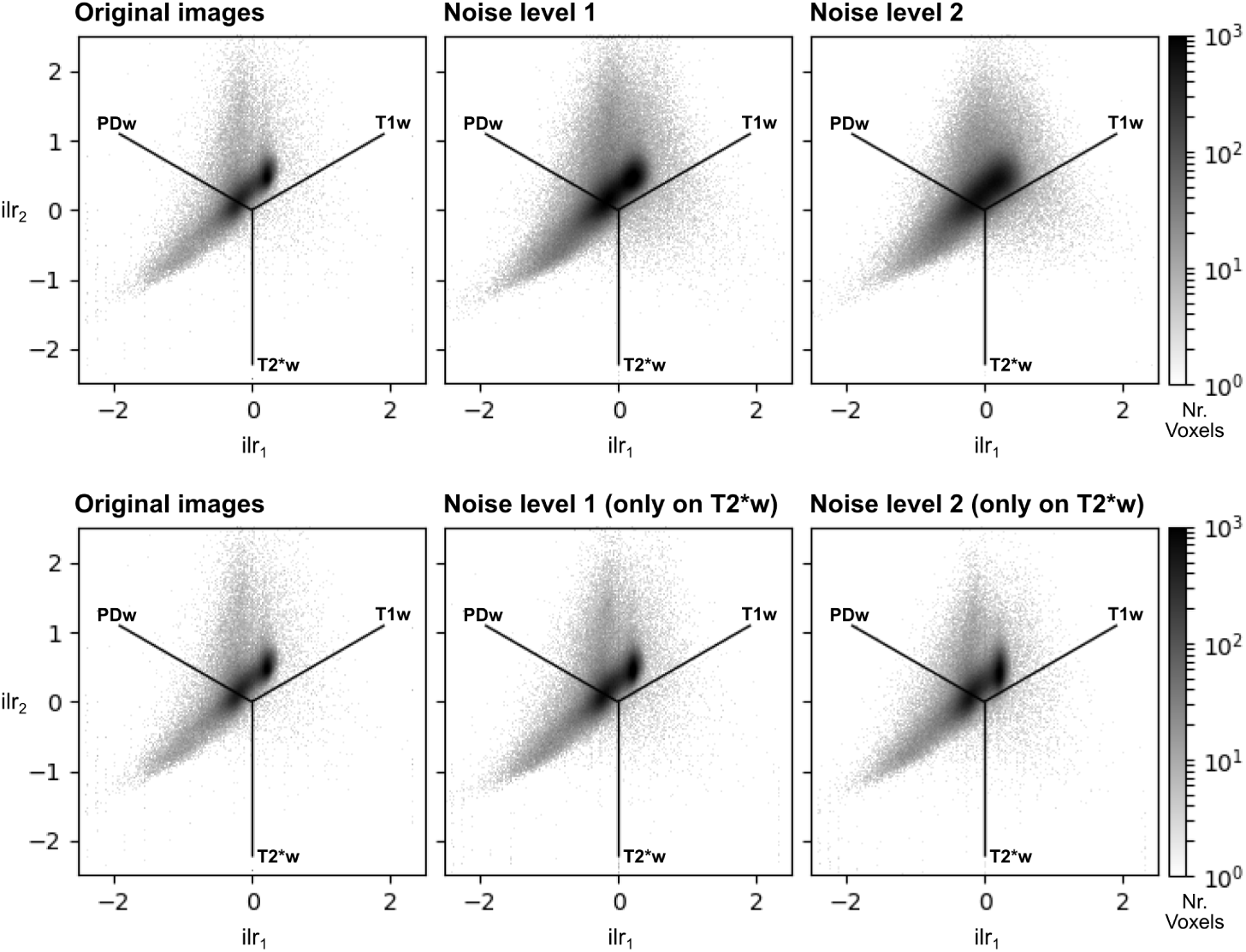
Impact of additional noise on CoDa method II. Shown are 2D histograms resulting from the CoDa method without any additional noise (left) and after applying a moderate (middle) and high (right column) amount of noise (see S3 Fig for additional details). Noise was either applied to all three channels equally (top row) or only to the T2*w image (bottom row). Noise causes structures in the 2D histogram that are initially well-defined to spread outward and, at very high noise levels, to lose shape. The histograms are based on data for one exemplary subject (sub-02).

**S5 Fig.**
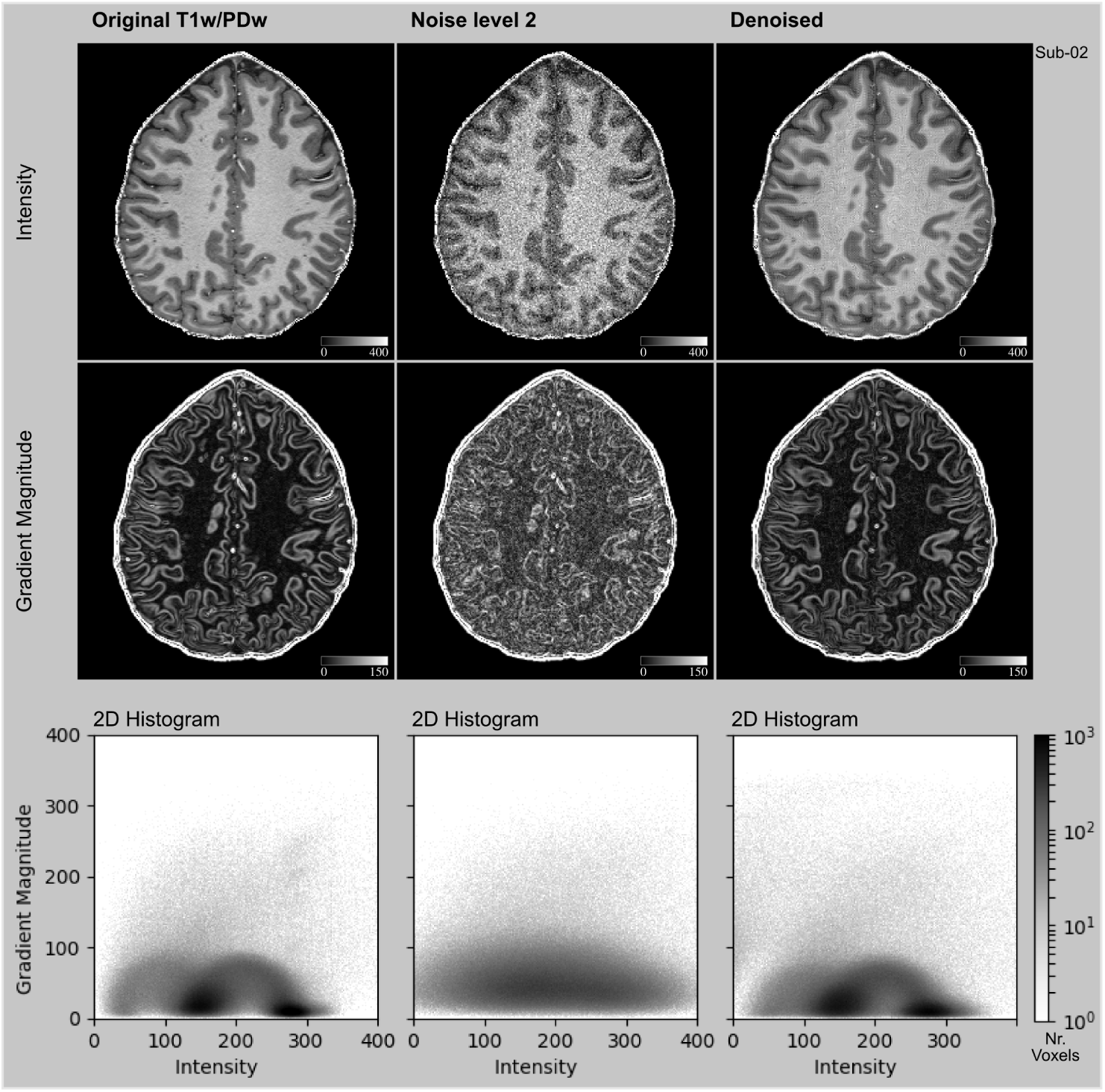
Noisy images can be de-noised using non-linear anisotropic smoothing. Shown are intensity images (top row), gradient magnitude images (middle row) and 2D histograms for the GraMag method (bottom row) for a T1w-divided-PDw MRI ratio image without any additional noise (left), after applying a high amount of noise (see S2 Fig for additional details) (middle), and after smoothing the noise-affected image (right column). As previously seen, noise causes structures in the 2D histogram to spread outward and to lose shape. This process can be reversed and noise-affected images can thus be recovered if an edge-preserving smoothing (see [66]) is applied. With smoothing, structures become more confined to the expected regions and well-defined shapes are regained. Images show a transverse slice for an exemplary subject (sub-02).

**S6 Fig.**
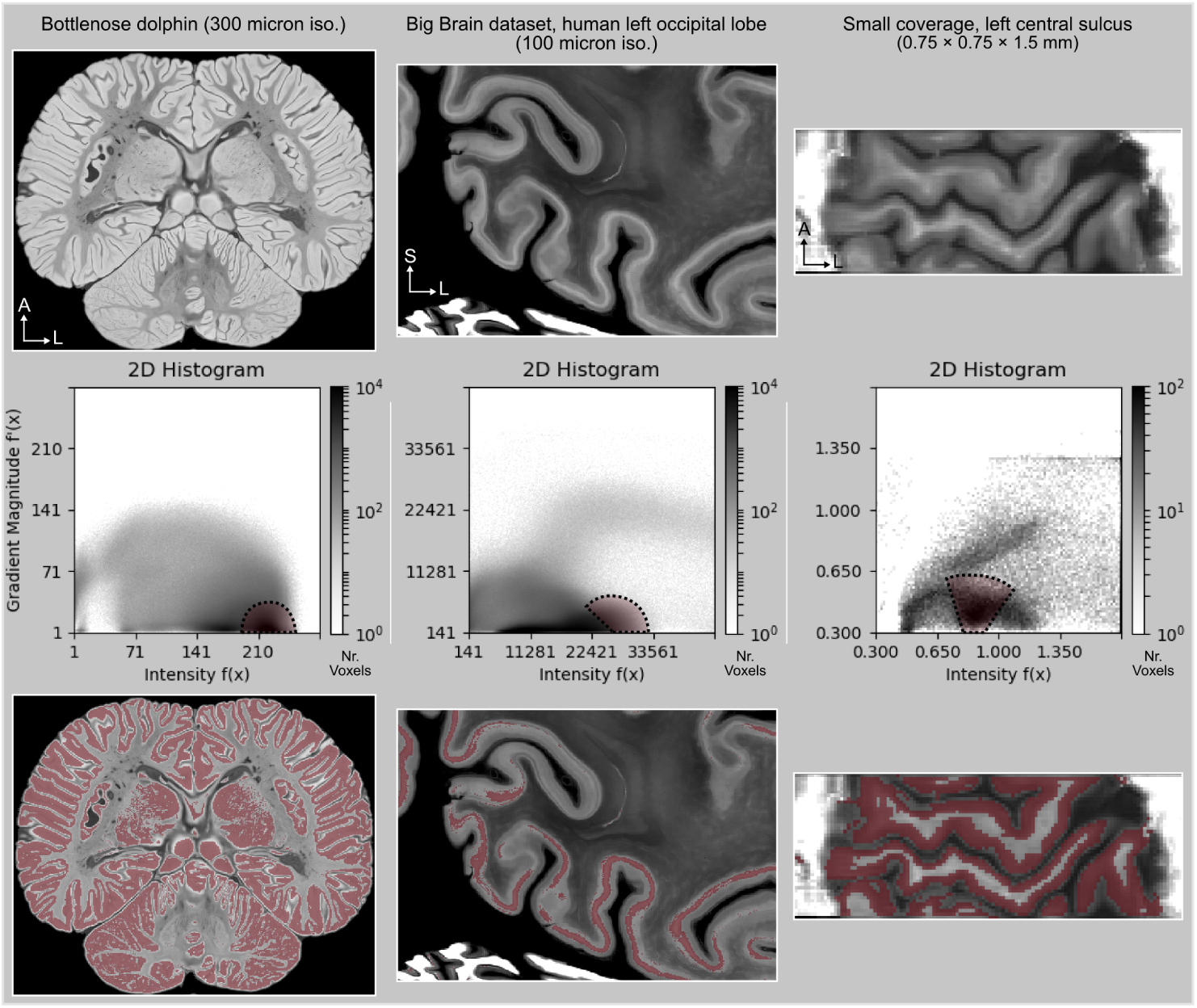
Application of GraMag to extra-ordinary MR images. Shown are several examples of the variety of existing volumetric datasets for which our methods appear to be useful. Every column represents different images: the brain of a bottle-nose dolphin [92] (left), the occipital lobe of a human brain with 100 micron resolution [93] (middle) and a human motor cortex acquired with small partial coverage (T1w EPI) with anisotropic resolution [94] (right). For every image we show a slice (top row), selected voxels in the 2D histogram (middle row) and selected voxels overlaid on the slice (bottom row). These images do not contain large intensity inhomogeneities. Therefore, no bias-field correction was performed. Mild non-linear anisotropic diffusion-based smoothing was applied to enhance CNR.

**S7 Fig.**
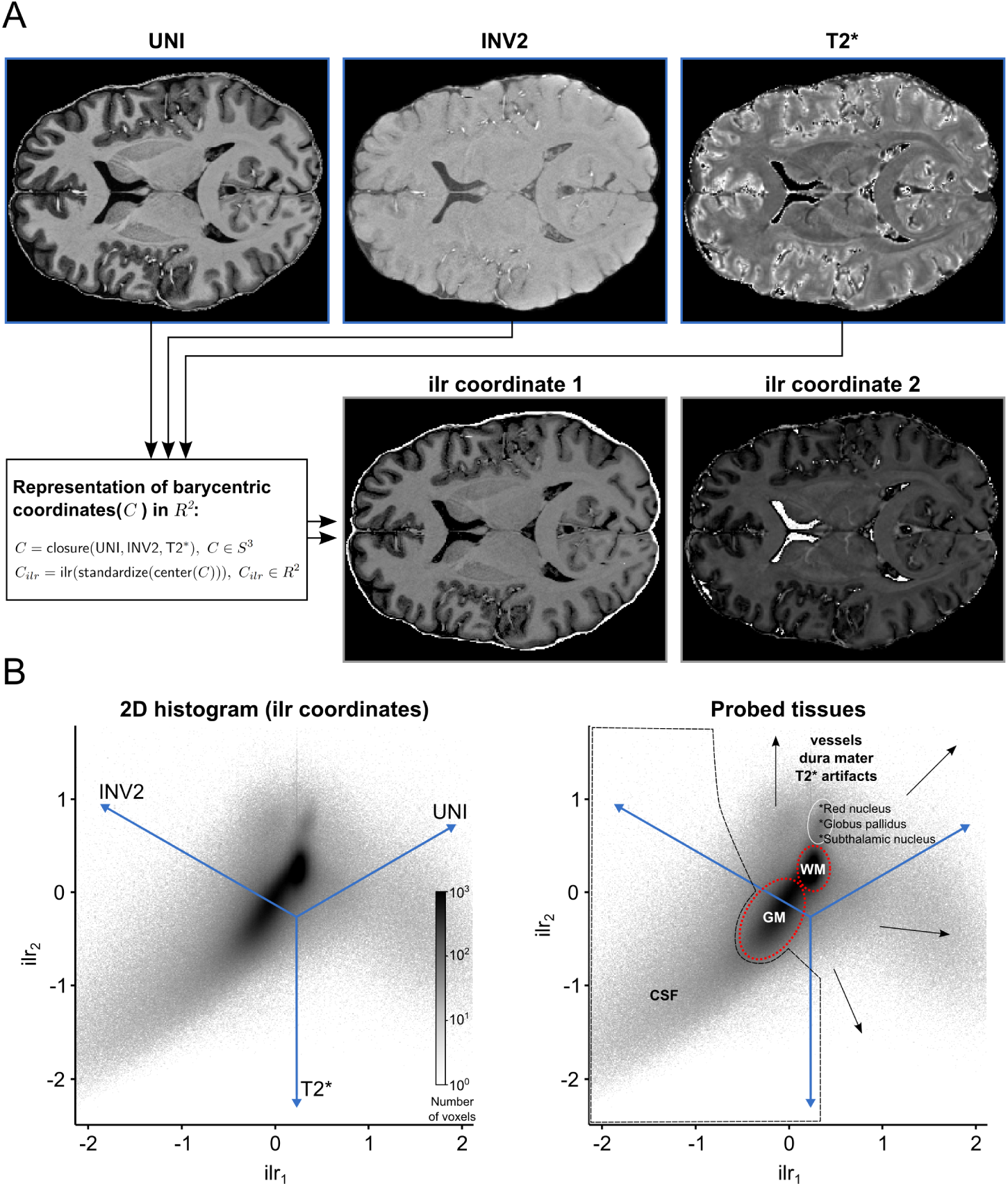
2D histogram representation of three 3D MRI contrast images. (A) Each voxel is considered as a three part composition. The barycentric coordinates of each composition which reside in 3D simplex space are represented in 2D real space after using a isometric log-ratio (ilr) transformation. (B) The ilr coordinates are used to create 2D histograms representing all voxels in the images. The blue lines are the embedded 3D real space primary axes (note that the input image units were initially normalized to have similar dynamic ranges to account for the large scale difference between T2* and MP2RAGE images). In this case, the ilr coordinates are not easily interpretable by themselves but they are useful to visualize the barycentric coordinates which are interpretable via the embedded real space axes. Darker regions in the histogram indicate that many voxels are characterized by this particular scale invariant combination of the image contrasts. In this representation, brain tissue (WM and GM, red dashed lines) becomes separable from non-brain tissue (black dashed lines and arrows). If desired, subcortical structures like the red nucleus, the globus pallidus and the subthalamic nucleus can additionally be identified (white circle).

**S8 Fig.**
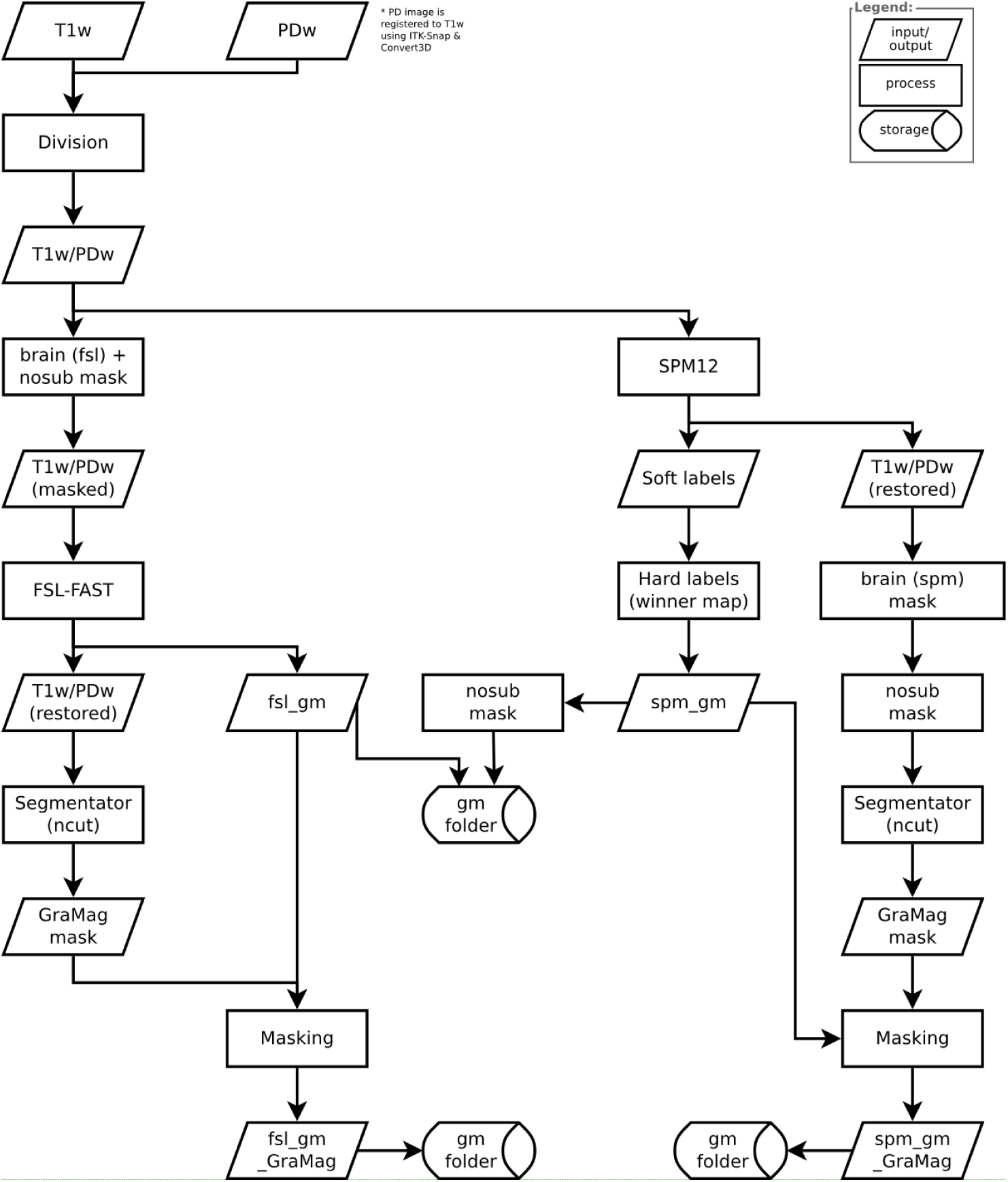
Flowchart diagram MPRAGE GraMag pipeline. This diagram provides a detailed overview of all the inputs, processing steps and outputs for MPRAGE GraMag pipeline. Rectangular shapes represent processing steps, rhombic shapes represent input or outputs and cylindrical shapes represent input or output locations.

**S9 Fig.**
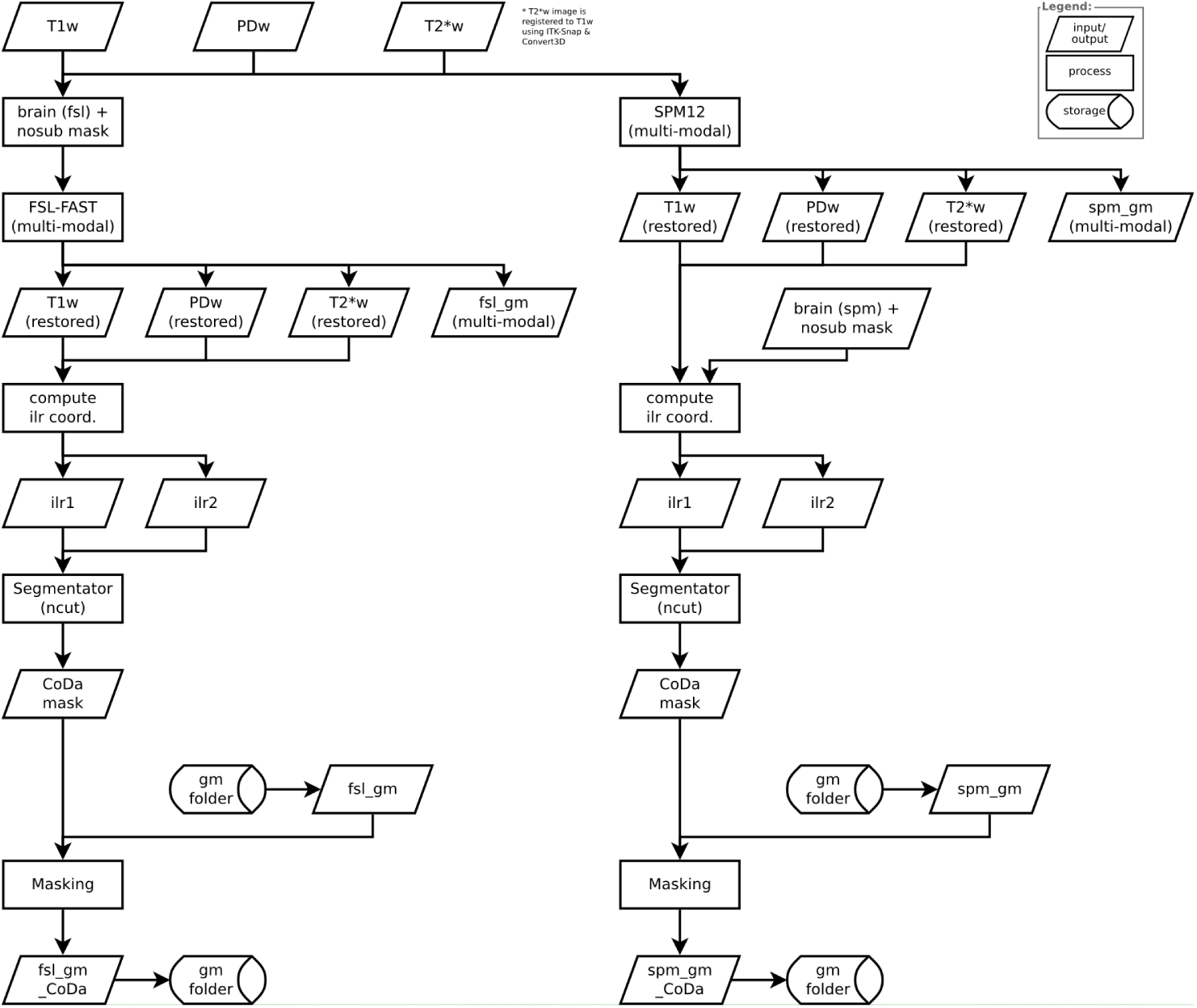
Flowchart diagram MPRAGE CoDa pipeline. This diagram provides a detailed overview of all the inputs, processing steps and outputs for MPRAGE CoDa pipeline. Rectangular shapes represent processing steps, rhombic shapes represent input or outputs and cylindrical shapes represent input or output locations.

**S10 Fig.**
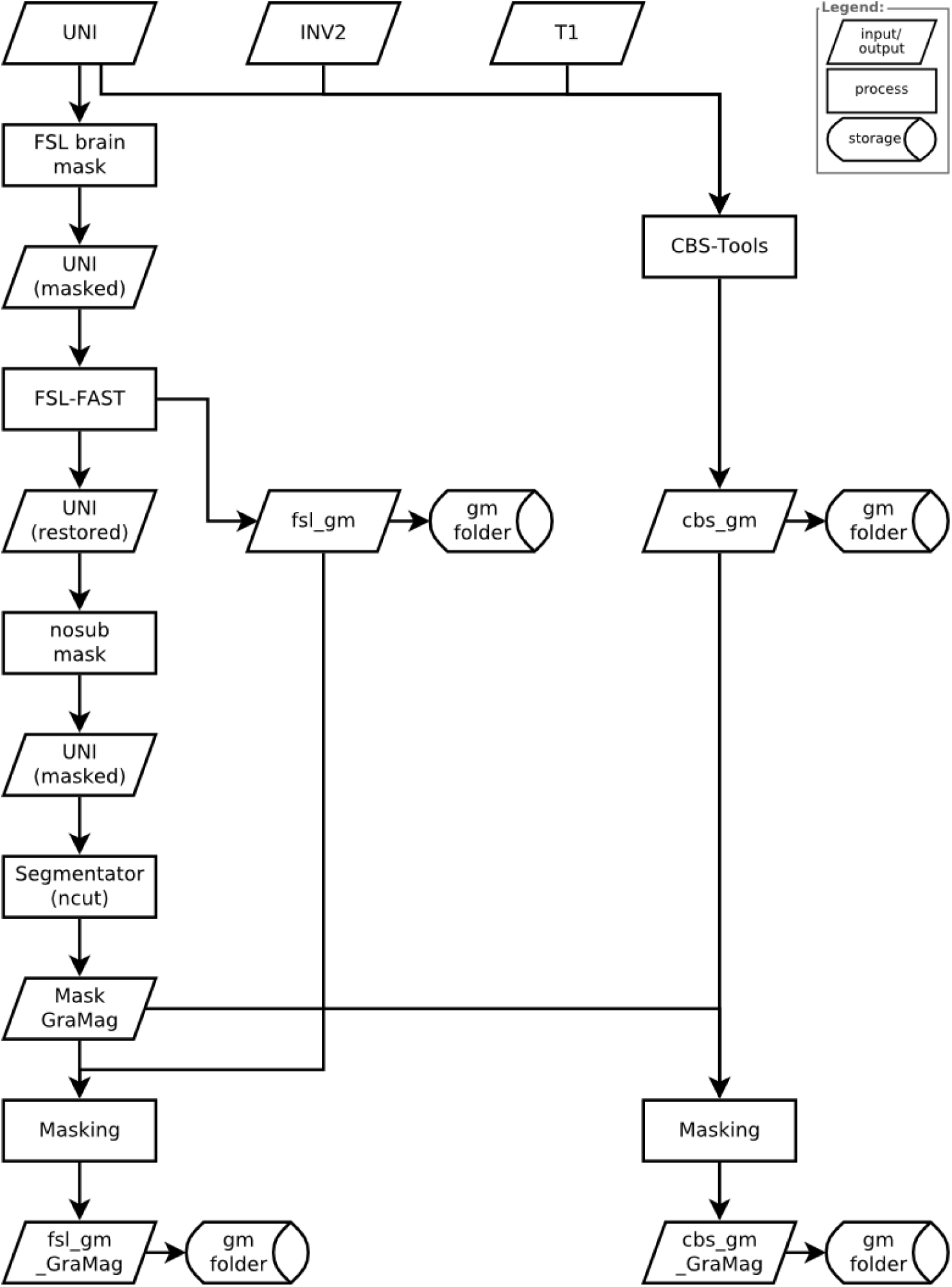
Flowchart diagram MP2RAGE GraMag pipeline. This diagram provides a detailed overview of all the inputs, processing steps and outputs for MP2RAGE GraMag pipeline. Rectangular shapes represent processing steps, rhombic shapes represent input or outputs and cylindrical shapes represent input or output locations.

**S11 Fig.**
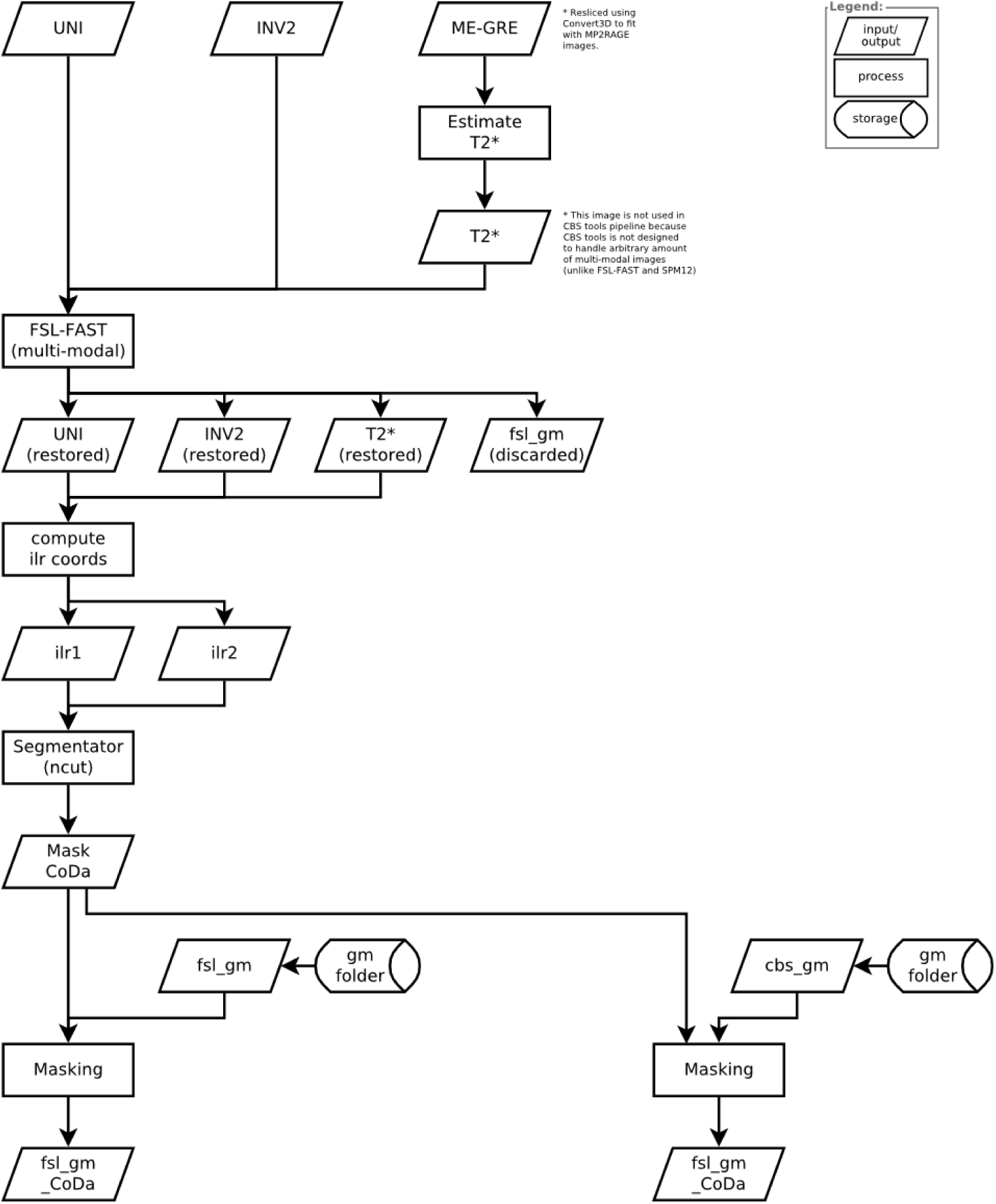
Flowchart diagram MP2RAGE CoDa pipeline. This diagram provides a detailed overview of all the inputs, processing steps and outputs for MP2RAGE CoDa pipeline. Rectangular shapes represent processing steps, rhombic shapes represent input or outputs and cylindrical shapes represent input or output locations.

**S12 Fig.**
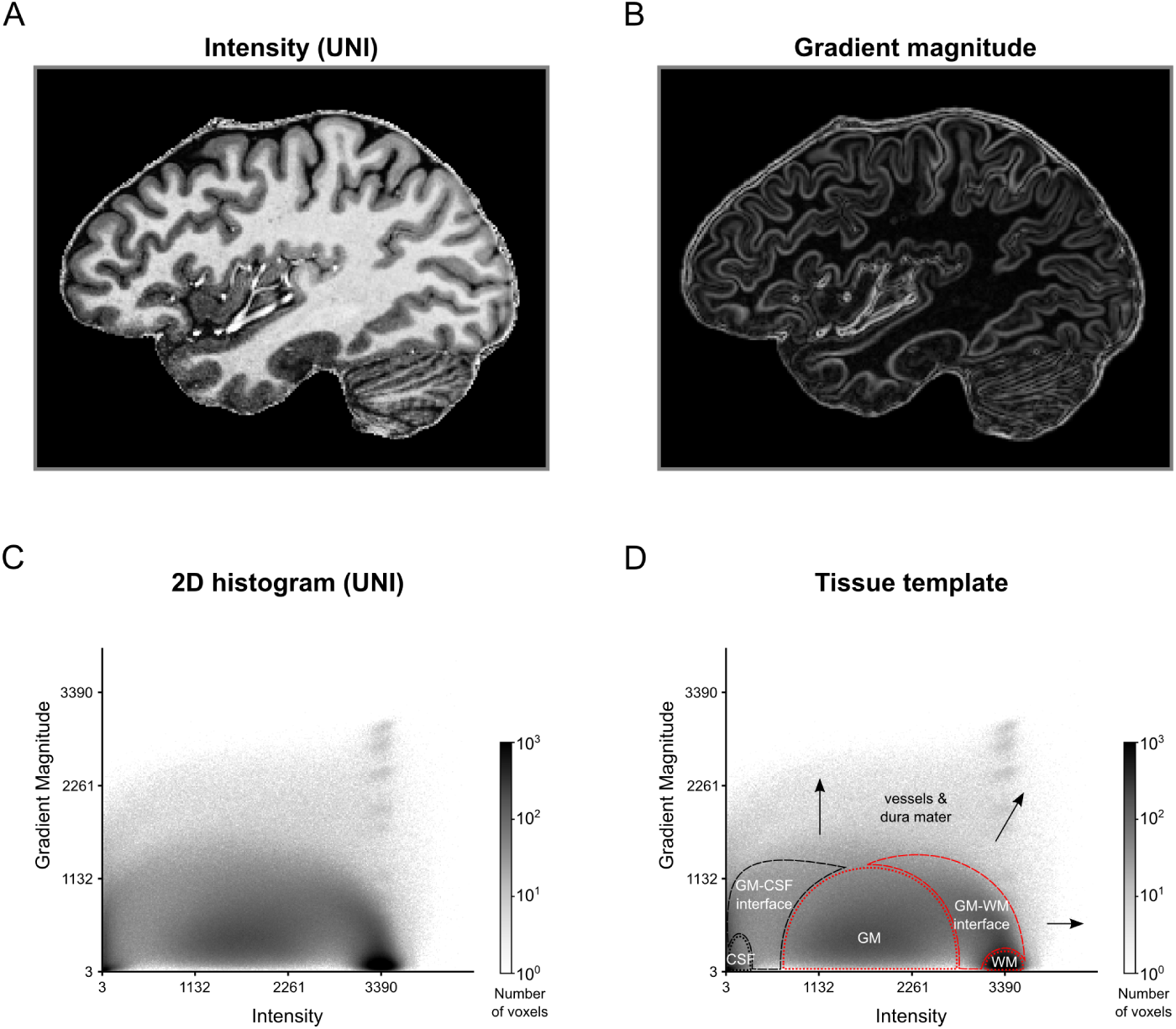
2D Histogram Representation for MRI Image of a Human Brain. The intensity (A) and gradient magnitude (B) values of a T1w-divided-by-PDw MRI image (MP2RAGE, 0.7 mm isotropic resolution) are represented in a 2D histogram (C). Darker regions in the histogram indicate that many voxels are characterized by this particular combination of image intensity and gradient magnitude. The 2D histogram displays a characteristic pattern with tissue types occupying particular areas of the histogram (D). Voxels containing CSF, dura mater or blood vessels (black dashed lines and arrows) cover different regions of the histogram than voxels containing WM and GM (red dashed lines). As a result, brain tissue becomes separable from non-brain tissue.

**S1 Table.** Segmentation performance MPRAGE data without artifact masking. The table shows the DICE (larger is better) and AVHD (less is better) for the initial SPM 12 and FSL FAST GM segmentations as well as after additional polishing, using either the gradient magnitude or the compositional data method.

|  |  | SPM |  | FAST |  |
| --- | --- | --- | --- | --- | --- |
|  |  | DICE | AVHD | DICE | AVHD |
| S02 |  |  |  |  |  |
|  | Init | 0.8576 | 0.7413 | 0.8896 | 0.5375 |
|  | Init + GraMag | 0.8594 | 0.5865 | 0.8902 | 0.4980 |
|  | Init + CoDa | 0.8525 | 0.6061 | 0.8421 | 0.5557 |
| S03 |  |  |  |  |  |
|  | Init | 0.8644 | 0.8753 | 0.8694 | 0.5739 |
|  | Init + GraMag | 0.8376 | 0.6139 | 0.8781 | 0.5612 |
|  | Init + CoDa | 0.8483 | 0.6299 | 0.8386 | 0.5687 |
| S05 |  |  |  |  |  |
|  | Init | 0.8569 | 0.6379 | 0.8782 | 0.5230 |
|  | Init + GraMag | 0.8688 | 0.5647 | 0.8909 | 0.5233 |
|  | Init + CoDa | 0.8596 | 0.5591 | 0.8692 | 0.5150 |
| S06 |  |  |  |  |  |
|  | Init | 0.8601 | 0.7789 | 0.8635 | 0.6477 |
|  | Init + GraMag | 0.8435 | 0.5797 | 0.8616 | 0.5909 |
|  | Init + CoDa | 0.8442 | 0.6287 | 0.8580 | 0.5530 |
| S07 |  |  |  |  |  |
|  | Init | 0.8897 | 0.6403 | 0.8490 | 0.7228 |
|  | Init + GraMag | 0.8893 | 0.5574 | 0.8460 | 0.6862 |
|  | Init + CoDa | 0.8508 | 0.5974 | 0.7935 | 0.7305 |

DICE, DICE Coefficient; AVHD, Average Hausdorff Distance; GraMag, gradient magnitude method; CoDa, compositional data method.

**S2 Table.**
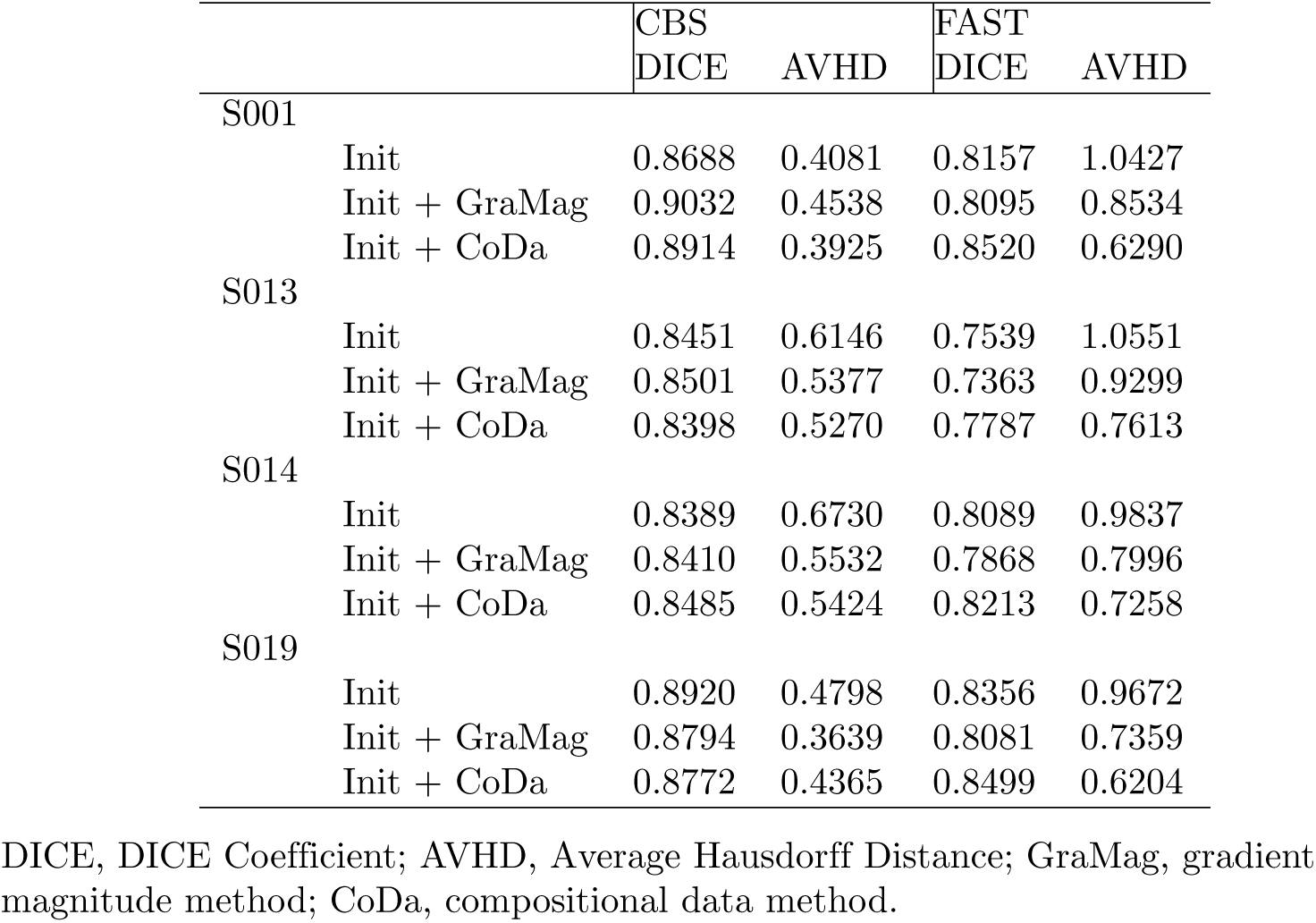
Segmentation performance MP2RAGE data without artifact masking. The table shows the DICE (larger is better) and AVHD (less is better) for the initial CBS tools and FSL FAST GM segmentations as well as after additional masking, using either the gradient magnitude or the compositional data method.

## References

1. Vaughan JT, Garwood M, Collins CM, Liu W, Delabarre L, Adriany G, et al. 7T vs. 4T: RF power, homogeneity, and signal-to-noise comparison in head images. Magnetic Resonance in Medicine. 2001;46(1):24–30. doi:10.1002/mrm.1156.

2. Duyn JH. The future of ultra-high field MRI and fMRI for study of the human brain. NeuroImage. 2012;62(2):1241–8. doi:10.1016/j.neuroimage.2011.10.065.

3. Ugurbil K. Magnetic resonance imaging at ultrahigh fields. IEEE Transactions on Biomedical Engineering. 2014;61(5):1364–1379. doi:10.1109/TBME.2014.2313619.

4. Martino FD, Yacoub E, Kemper V, Moerel M, Uludag K, Weerd PD, et al. The impact of ultra-high field MRI on cognitive and computational neuroimaging. NeuroImage. 2017;doi:https://doi.org/10.1016/j.neuroimage.2017.03.060.

5. Kemper VG, Martino FD, Emmerling TC, Yacoub E, Goebel R. High resolution data analysis strategies for mesoscale human functional MRI at 7 and 9.4T. NeuroImage. 2017;doi:https://doi.org/10.1016/j.neuroimage.2017.03.058.

6. Dumoulin SO, Fracasso A, van der Zwaag W, Siero JCW, Petridou N. Ultra-high field MRI: Advancing systems neuroscience towards mesoscopic human brain function. NeuroImage. 2017;doi:https://doi.org/10.1016/j.neuroimage.2017.01.028.

7. Polimeni JR, Renvall V, Zaretskaya N, Fischl B. Analysis strategies for high-resolution UHF-fMRI data. NeuroImage. 2017;doi:https://doi.org/10.1016/j.neuroimage.2017.04.053.

8. Trampel R, Bazin PL, Pine K, Weiskopf N. In-vivo magnetic resonance imaging (MRI) of laminae in the human cortex. NeuroImage. 2017;(September):1–9. doi:10.1016/j.neuroimage.2017.09.037.

9. Polimeni JR, Fischl B, Greve DN, Wald LL. Laminar analysis of 7T BOLD using an imposed spatial activation pattern in human V1. NeuroImage. 2010;52(4):1334–1346. doi:10.1016/j.neuroimage.2010.05.005.

10. Koopmans PJ, Barth M, Orzada S, Norris DG. Multi-echo fMRI of the cortical laminae in humans at 7 T. NeuroImage. 2011;56(3):1276–1285. doi:10.1016/j.neuroimage.2011.02.042.

11. De Martino F, Zimmermann J, Muckli L, Ugurbil K, Yacoub E, Goebel R. Cortical Depth Dependent Functional Responses in Humans at 7T: Improved Specificity with 3D GRASE. PLoS ONE. 2013;8(3):e60514. doi:10.1371/journal.pone.0060514.

12. Huber L, Goense J, Kennerley AJ, Trampel R, Guidi M, Reimer E, et al. Cortical lamina-dependent blood volume changes in human brain at 7 T. NeuroImage. 2015;107:23–33. doi:10.1016/j.neuroimage.2014.11.046.

13. Muckli L, De Martino F, Vizioli L, Petro LS, Smith FW, Ugurbil K, et al. Contextual Feedback to Superficial Layers of V1. Current Biology. 2015;25(20):2690–2695. doi:10.1016/j.cub.2015.08.057.

14. Kok P, Bains LJ, Van Mourik T, Norris DG, De Lange FP. Selective activation of the deep layers of the human primary visual cortex by top-down feedback. Current Biology. 2016;26(3):371–376. doi:10.1016/j.cub.2015.12.038.

15. Waehnert MD, Dinse J, Weiss M, Streicher MN, Waehnert P, Geyer S, et al. Anatomically motivated modeling of cortical laminae. NeuroImage. 2014;93:210–220. doi:10.1016/j.neuroimage.2013.03.078.

16. Yacoub E, Harel N, Ugurbil K. High-field fMRI unveils orientation columns in humans. Proceedings of the National Academy of Sciences. 2008;105(30):10607–10612. doi:10.1073/pnas.0804110105.

17. Zimmermann J, Goebel R, de Martino F, van de Moortele PF, Feinberg D, Adriany G, et al. Mapping the organization of axis of motion selective features in human area mt using high-field fmri. PLoS ONE. 2011;6(12):1–10. doi:10.1371/journal.pone.0028716.

18. De Martino F, Moerel M, Ugurbil K, Goebel R, Yacoub E, Formisano E. Frequency preference and attention effects across cortical depths in the human primary auditory cortex. Proceedings of the National Academy of Sciences. 2015;112(52):16036–16041. doi:10.1073/pnas.1507552112.

19. Goncalves NR, Ban H, Sanchez-Panchuelo RM, Francis ST, Schluppeck D, Welchman AE. 7 Tesla fMRI Reveals Systematic Functional Organization for Binocular Disparity in Dorsal Visual Cortex. Journal of Neuroscience. 2015;35(7):3056–3072. doi:10.1523/JNEUROSCI.3047-14.2015.

20. Nasr S, Polimeni JR, Tootell RBH. Interdigitated Color- and Disparity-Selective Columns within Human Visual Cortical Areas V2 and V3. Journal of Neuroscience. 2016;36(6). doi:10.1523/JNEUROSCI.3518-15.2016.

21. Tootell RBH, Nasr S. Columnar Segregation of Magnocellular and Parvocellular Streams in Human Extrastriate Cortex. The Journal of Neuroscience. 2017; p. 0690–17. doi:10.1523/JNEUROSCI.0690-17.2017.

22. Harvey BM, Klein BP, Petridou N, Dumoulin SO. Topographic Representation of Numerosity in the Human Parietal Cortex. Science. 2013;341(6150):1123–1126. doi:10.1126/science.1239052.

23. Harvey BM, Fracasso A, Petridou N, Dumoulin SO. Topographic representations of object size and relationships with numerosity reveal generalized quantity processing in human parietal cortex. Proceedings of the National Academy of Sciences of the United States of America. 2015;112(44):13525–30. doi:10.1073/pnas.1515414112.

24. Fracasso A, Petridou N, Dumoulin SO. Systematic variation of population receptive field properties across cortical depth in human visual cortex. NeuroImage. 2016;139:427–438. doi:10.1016/j.neuroimage.2016.06.048.

25. Van de Moortele PF, Auerbach EJ, Olman C, Yacoub E, UȈgurbil K, Moeller S. T1 weighted brain images at 7 Tesla unbiased for Proton Density, T2* contrast and RF coil receive B1 sensitivity with simultaneous vessel visualization. NeuroImage. 2009;46(2):432–446. doi:10.1016/j.neuroimage.2009.02.009.

26. Marques JP, Kober T, Krueger G, van der Zwaag W, Van de Moortele PF, Gruetter R. MP2RAGE, a self bias-field corrected sequence for improved segmentation and T1-mapping at high field. NeuroImage. 2010;49(2):1271–1281. doi:10.1016/j.neuroimage.2009.10.002.

27. Viviani R, Pracht ED, Brenner D, Beschoner P, Stingl JC, Stöcker T. Multimodal MEMPRAGE, FLAIR, and R2* segmentation to resolve dura and vessels from cortical gray matter. Frontiers in Neuroscience. 2017;11(MAY):1–13. doi:10.3389/fnins.2017.00258.

28. van der Kouwe AJW, Benner T, Salat DH, Fischl B. Brain morphometry with multiecho MPRAGE. NeuroImage. 2008;40(2):559–569. doi:10.1016/j.neuroimage.2007.12.025.

29. Viviani R. A Digital Atlas of Middle to Large Brain Vessels and Their Relation to Cortical and Subcortical Structures. Frontiers in Neuroanatomy. 2016;10(February):12. doi:10.3389/FNANA.2016.00012.

30. Helms G, Kallenberg K, Dechent P. Contrast-driven approach to intracranial segmentation using a combination of T2- and T1-weighted 3D MRI data sets. Journal of Magnetic Resonance Imaging. 2006;24(4):790–795. doi:10.1002/jmri.20692.

31. Helms G. Segmentation of human brain using structural MRI. Magnetic Resonance Materials in Physics, Biology and Medicine. 2016;29(2):111–124. doi:10.1007/s10334-015-0518-z.

32. Bazin PL, Weiss M, Dinse J, Schafer A, Trampel R, Turner R. A computational framework for ultra-high resolution cortical segmentation at 7 Tesla. NeuroImage. 2014;93:201–209. doi:10.1016/j.neuroimage.2013.03.077.

33. Despotović I, Goossens B, Philips W. MRI segmentation of the human brain: challenges, methods, and applications. Computational and mathematical methods in medicine. 2015;2015:450341. doi:10.1155/2015/450341.

34. Renvall V, Witzel T, Wald LL, Polimeni JR. Automatic cortical surface reconstruction of high-resolution T1 echo planar imaging data. NeuroImage. 2016;134:338–354. doi:10.1016/j.neuroimage.2016.04.004.

35. Kashyap S, Ivanov D, Havlicek M, Poser BA, Uludağ K. Impact of acquisition and analysis strategies on cortical depth-dependent fMRI; 2017.

36. Kindlmann G, Durkin JW. Semi-automatic generation of transfer functions for direct volume rendering. In: Proceedings of the 1998 IEEE symposium on Volume visualization - VVS ’98. New York, New York, USA: ACM Press; 1998. p. 79–86.

37. Kniss J, Kindlmann G, Hansen C. Interactive volume rendering using multi-dimensional transfer functions and direct manipulation widgets. In: Proceedings Visualization, 2001. VIS ’01. IEEE; 2001. p. 255–562.

38. Kniss J, Kindlmann G, Hansen CD. Multidimensional transfer functions for volume rendering. Visualization Handbook. 2005; p. 189–209. doi:10.1016/B978-012387582-2/50011-3.

39. Ip CY, Varshney A, Jaja J. Hierarchical exploration of volumes using multilevel segmentation of the intensity-gradient histograms. IEEE Transactions on Visualization and Computer Graphics. 2012;18(12):2355–2363. doi:10.1109/TVCG.2012.231.

40. Jolliffe IT. Principal Component Analysis and Factor Analysis. In: Technometrics. vol. 30; 1986. p. 115–128. Available from: http://onlinelibrary.wiley.com/doi/10.1002/0470013192.bsa501/fullhttp://link.springer.com/10.1007/978-1-4757-1904-8{_}7.

41. Borg I, Groenen P. Modern Multidimensional Scaling: Theory and Applications. Chapter 10. 2005; p. 100–131.

42. Van Der Maaten L, Hinton G. Visualizing Data using t-SNE. Journal of Machine Learning Research. 2008;9:2579–2605. doi:10.1007/s10479-011-0841-3.

43. Venkataraju KU, Paiva ARC, Jurrus E, Tasdizen T. Automatic markup of neural cell membranes using boosted decision stumps. In: Proceedings - 2009 IEEE International Symposium on Biomedical Imaging: From Nano to Macro, ISBI 2009; 2009. p. 1039–1042.

44. Jain V, Turaga SC, Briggman K, Helmstaedter MN, Denk W, Seung HS. Learning to Agglomerate Superpixel Hierarchies. Advances in Neural Information Processing Systems. 2011; p. 648–656.

45. Liu T, Jurrus E. Watershed merge tree classification for electron microscopy image segmentation. … (ICPR), 2012 21st …. 2012;(Icpr):3–6. doi:10.1097/MPG.0b013e3181a15ae8.Screening.

46. Nunez-Iglesias J, Kennedy R, Parag T, Shi J, Chklovskii DB. Machine Learning of Hierarchical Clustering to Segment 2D and 3D Images. PLoS ONE. 2013;8(8). doi:10.1371/journal.pone.0071715.

47. Pawlowsky-Glahn V, Egozcue JJ, Tolosana-Delgado R. Modelling and Analysis of Compositional Data. Chichester, UK: John Wiley & Sons, Ltd; 2015.

48. Gulban OF, Schneider M. Segmentator v1.5.0; 2017. Available from: https://doi.org/10.5281/zenodo.1219243.

49. Gulban OF, Schneider M, Marquardt I, Haast RAM, De Martino F. Dataset: A scalable method to improve gray matter segmentation at ultra high field MRI.; 2017. Available from: https://doi.org/10.5281/zenodo.1117859.

50. Schneider M, Gulban OF. Processing scripts: A scalable method to improve gray matter segmentation at ultra high field MRI.; 2018. Available from: https://doi.org/10.5281/zenodo.1217084.

51. Kniss J, Kindlmann G, Hansen C. Multidimensional transfer functions for interactive volume rendering. IEEE Transactions on Visualization and Computer Graphics. 2002;8(3):270–285. doi:10.1109/TVCG.2002.1021579.

52. Ljung P, Krüger J, Groller E, Hadwiger M, Hansen CD, Ynnerman A. State of the Art in Transfer Functions for Direct Volume Rendering. Computer Graphics Forum. 2016;35(3):669–691. doi:10.1111/cgf.12934.

53. Glasser MF, Van Essen DC. Mapping Human Cortical Areas In Vivo Based on Myelin Content as Revealed by T1- and T2-Weighted MRI. Journal of Neuroscience. 2011;31(32):11597–11616. doi:10.1523/JNEUROSCI.2180-11.2011.

54. De Martino F, Moerel M, Xu J, Van De Moortele PF, Ugurbil K, Goebel R, et al. High-resolution mapping of myeloarchitecture in vivo: Localization of auditory areas in the human brain. Cerebral Cortex. 2015;25(10):3394–3405. doi:10.1093/cercor/bhu150.

55. Fauvel J, Wilson R, Flood R. Möbius and his Band: Mathematics and astronomy in nineteenth-century Germany. Oxford, England: Oxford University Press; 1993.

56. Munkres JR. Elements of Algebraic Topology. 1st ed. Addison-Wesley; 1984.

57. Aitchison J. The Statistical Analysis of Compositional Data. Journal of the Royal Statistical Society. 1982;44(2):139–177.

58. Aitchison J. A Concise Guide to Compositional Data Analysis. CDA Workshop Girona. 2002;24:73–81.

59. Egozcue JJ, Pawlowsky-Glahn V, Mateu-Figueras G, Barceló-Vidal C. Isometric Logratio Transformations for Compositional Data Analysis. Mathematical Geology. 2003;35(3):279–300. doi:10.1023/A:1023818214614.

60. Lancaster HO. The Helmert Matrices. The American Mathematical Monthly. 1965;72(1):4. doi:10.2307/2312989.

61. Tsagris MT, Preston S, Wood ATA. A data-based power transformation for compositional data. arXiv preprint. 2011;(1):1–9.

62. Smith SM. Fast robust automated brain extraction. Human Brain Mapping. 2002;17(3):143–155. doi:10.1002/hbm.10062.

63. Ashburner J, Friston KJ. Unified segmentation. NeuroImage. 2005;26(3):839–851. doi:10.1016/j.neuroimage.2005.02.018.

64. Lüsebrink F, Sciarra A, Mattern H, Yakupov R, Speck O. Data from: T1-weighted in vivo human whole brain MRI dataset with an ultrahigh isotropic resolution of 250 µm; 2017. Available from: https://doi.org/10.5061/dryad.38s74.

65. Lusebrink F, Sciarra A, Mattern H, Yakupov R, Speck O. T1-weighted in vivo human whole brain MRI dataset with an ultrahigh isotropic resolution of 250 µm. Scientific Data. 2017;4:1–11. doi:10.1038/sdata.2017.32.

66. Weickert J. Anisotropic diffusion in image processing. Image Rochester NY. 1998;256(3):170. doi:10.1.1.11.751.

67. Mirebeau J, Fehrenbach J, Risser L, Tobji S. Anisotropic Diffusion in ITK. CoRR. 2015;abs/1503.00992.

68. Teeuwisse WM, Brink WM, Webb AG. Quantitative assessment of the effects of high-permittivity pads in 7 tesla MRI of the brain. Magnetic resonance in medicine. 2012;67(5):1285–1293.

69. Griswold MA, Jakob PM, Heidemann RM, Nittka M, Jellus V, Wang J, et al. Generalized Autocalibrating Partially Parallel Acquisitions (GRAPPA). Magnetic Resonance in Medicine. 2002;47(6):1202–1210. doi:10.1002/mrm.10171.

70. Haast RAM, Ivanov D, Formisano E, Uludağ K. Reproducibility and Reliability of Quantitative and Weighted T1 and T2(*) Mapping for Myelin-Based Cortical Parcellation at 7 Tesla. Frontiers in neuroanatomy. 2016;10:112. doi:10.3389/fnana.2016.00112.

71. Yushkevich PA, Piven J, Cody Hazlett H, Gimpel Smith R, Ho S, Gee JC, et al. User-Guided 3D Active Contour Segmentation of Anatomical Structures: Significantly Improved Efficiency and Reliability. Neuroimage. 2006;31(3):1116–1128.

72. Van Der Walt S, Colbert SC, Varoquaux G. The NumPy array: a structure for efficient numerical computation. Computing in Science & Engineering. 2011;13(2):22–30.

73. Jones E, Oliphant T, Peterson P, et al. SciPy: Open source scientific tools for Python; 2001–. Available from: http://www.scipy.org/.

74. Hunter JD. Matplotlib: A 2D graphics environment. Computing In Science & Engineering. 2007;9(3):90–95. doi:10.1109/MCSE.2007.55.

75. Brett M, Hanke M, Côté MA, Markiewicz C, Ghosh S, Wassermann D, et al. nipy/nibabel: 2.2.0; 2017. Available from: https://doi.org/10.5281/zenodo.1011207.

76. van der Walt S, Schönberger JL, Nunez-Iglesias J, Boulogne F, Warner JD, Yager N, et al. scikit-image: image processing in Python. PeerJ. 2014;2:e453. doi:10.7717/peerj.453.

77. Scharr H. Optimale operatoren in der digitalen bildverarbeitung; 2000.

78. Jähne B, Haußecker H. Handbook of computer vision and applications. vol. 2. Elsevier; 2000.

79. Gulban OF. Compoda v0.3.3; 2018. Available from: https://doi.org/10.5281/zenodo.1209137.

80. Zhang Y, Brady M, Smith S. Segmentation of brain MR images through a hidden Markov random field model and the expectation-maximization algorithm. IEEE Transactions on Medical Imaging. 2001;20(1):45–57. doi:10.1109/42.906424.

81. Taha AA, Hanbury A. Metrics for evaluating 3D medical image segmentation: analysis, selection, and tool. BMC Medical Imaging. 2015;15(1):29. doi:10.1186/s12880-015-0068-x.

82. Sereno MI, Lutti A, Weiskopf N, Dick F. Mapping the human cortical surface by combining quantitative T1 with retinotopy. Cerebral Cortex. 2013;23(9):2261–2268. doi:10.1093/cercor/bhs213.

83. Dick F, Taylor Tierney A, Lutti A, Josephs O, Sereno MI, Weiskopf N. In Vivo Functional and Myeloarchitectonic Mapping of Human Primary Auditory Areas. Journal of Neuroscience. 2012;32(46):16095–16105. doi:10.1523/JNEUROSCI.1712-12.2012.

84. Marques JP, Gruetter R. New Developments and Applications of the MP2RAGE Sequence - Focusing the Contrast and High Spatial Resolution R1 Mapping. PLoS ONE. 2013;8(7). doi:10.1371/journal.pone.0069294.

85. Gulban OF. The relation between color spaces and compositional data analysis demonstrated with magnetic resonance image processing applications; 2017. Available from: https://arxiv.org/abs/1705.03457.

86. Collins DL, Zijdenbos aP, Kollokian V, Sled JG, Kabani NJ, Holmes CJ, et al. Design and construction of a realistic digital brain phantom. IEEE Transactions on Medical Imaging. 1998;17(3):463–468. doi:10.1109/42.712135.

87. Aubert-Broche B, Evans AC, Collins L. A new improved version of the realistic digital brain phantom. NeuroImage. 2006;32(1):138–145. doi:https://doi.org/10.1016/j.neuroimage.2006.03.052.

88. Aubert-Broche B, Griffin M, Pike GB, Evans AC, Collins DL. Twenty New Digital Brain Phantoms for Creation of Validation Image Data Bases. IEEE Trans Med Imaging. 2006;25(11):1410–1416.

89. Valverde S, Oliver A, Cabezas M, Roura E, Lladó X. Comparison of 10 brain tissue segmentation methods using revisited IBSR annotations. Journal of Magnetic Resonance Imaging. 2015;41(1):93–101. doi:10.1002/jmri.24517.

90. Sequence-independent segmentation of magnetic resonance images. In: NeuroImage. vol. 23; 2004.

91. Goebel R, Esposito F, Formisano E. Analysis of Functional Image Analysis Contest (FIAC) data with BrainVoyager QX: From single-subject to cortically aligned group General Linear Model analysis and self-organizing group Independent Component Analysis. Human Brain Mapping. 2006;27(5):392–401. doi:10.1002/hbm.20249.

92. Toro R, Grisanti F, Herbin M, Santin M. The Brain Catalogue: An open portal for comparative neuroanatomy; 2014. Available from: https://figshare.com/articles/The_Brain_Catalogue_An_open_portal_for_comparative_neuroanatomy/1048827/1.

93. Amunts K, Lepage C, Borgeat L, Mohlberg H, Dickscheid T, Rousseau MÉ, et al. BigBrain: An ultrahigh-resolution 3D human brain model. Science. 2013;340(6139):1472–1475. doi:10.1126/science.1235381.

94. Huber L, Handwerker DA, Jangraw DC, Chen G, Hall A, Stüber C, et al. High-Resolution CBV-fMRI Allows Mapping of Laminar Activity and Connectivity of Cortical Input and Output in Human M1. Neuron. 2017;96(6):1253–1263.e7. doi:10.1016/j.neuron.2017.11.005.

